# An Ancestral Balanced Inversion Polymorphism Confers Global Adaptation

**DOI:** 10.1101/2023.01.31.526462

**Authors:** Martin Kapun, Esra Durmaz Mitchell, Tadeusz J. Kawecki, Paul Schmidt, Thomas Flatt

## Abstract

Since the pioneering work of Dobzhansky in the 1930s and 1940s, many chromosomal inversions have been identified but how they contribute to adaptation remains poorly understood. In *Drosophila melanogaster*, the widespread inversion polymorphism *In(3R)Payne* underpins latitudinal clines in fitness traits on multiple continents. Here, we use single-individual whole-genome sequencing, transcriptomics and published sequencing data to study the population genomics of this inversion on four continents: in its ancestral African range and in derived populations in Europe, North America, and Australia. Our results confirm that this inversion originated in sub-Saharan Africa and subsequently became cosmopolitan; we observe marked monophyletic divergence of inverted and non-inverted karyotypes, with some substructure among inverted chromosomes between continents. Despite divergent evolution of this inversion since its out-of-Africa migration, derived non-African populations exhibit similar patterns of long-range linkage disequilibrium between the inversion breakpoints and major peaks of divergence in its center, consistent with balancing selection and suggesting that the inversion harbors alleles that are maintained by selection on several continents. Using RNA-seq we identify overlap between inversion-linked SNPs and loci that are differentially expressed between inverted and non-inverted chromosomes. Expression levels are higher for inverted chromosomes at low temperature, suggesting loss of buffering or compensatory plasticity and consistent with higher inversion frequency in warm climates. Our results suggest that this ancestrally tropical balanced polymorphism spread around the world and became latitudinally assorted along similar but independent climatic gradients, always being frequent in subtropical/tropical areas but rare or absent in temperate climates.

## Introduction

Chromosomal inversions are structural mutations that cause the gene order of a chromosomal segment to be reversed (Sturtevant 1917, 1919, 1921). Because inversions suppress crossing-over (but not gene conversion events) in heterozygous state, they can cause an effective barrier to genetic exchange (‘gene flux’) between inverted and non-inverted (‘standard’) chromosomes (Rozas and Aguadé 1994; Navarro et al. 1997; Griffiths et al. 2000; Schaeffer and Anderson 2005; Kirkpatrick 2010; Charlesworth 2016; Crown et al. 2018; Korunes and Noor 2019; Kapun and Flatt 2019; Durmaz et al. 2020). This pervasive effect of inversions on patterns of recombination can have major evolutionary consequences. For example, inversions can contribute to (i) speciation by allowing mutations involved in reproductive isolation to accumulate; (ii) genetic divergence between the sexes by accumulating on sex chromosomes; and (iii) adaptation by capturing beneficial alleles at multiple loci and binding them together (Dobzhansky 1948, 1949, 1950; Charlesworth and Charlesworth 1973; Rieseberg 2001; Noor et al. 2001; Navarro and Barton 2003; Kirkpatrick and Barton 2006; Hoffmann and Rieseberg 2008; Kirkpatrick 2010; Charlesworth 2016; Fuller et al. 2016, 2017; Charlesworth and Barton 2018; Wellenreuther and Bernatchez 2018; Faria et al. 2019; Fuller et al. 2019; Kapun and Flatt 2019; Durmaz et al. 2020; Charlesworth and Flatt 2021; Mackintosh et al. 2022).

Since the discovery of inversions in the early 20^th^ century by Sturtevant (1917, 1919, 1921), their role in adaptation has attracted great interest among evolutionary geneticists (Dobzhansky 1948, 1950; Krimbas and Powell 1992; Hoffmann et al. 2004; Kirkpatrick and Barton 2006; Hoffmann and Rieseberg 2008; Kirkpatrick 2010; Guerrero et al. 2012; Wellenreuther and Bernatchez 2018; Faria et al. 2019; Kapun and Flatt 2019). For instance, theory suggests that linked selection can cause the spread of an initially rare inversion when it captures a locally adaptive haplotype, protects it from recombination load and/or maladaptive gene flow from neighboring populations, and then ‘hitchhikes’ with it to high frequency; alternatively, a new inversion might be favored by direct positive selection when the breakpoints of the inversion fortuitously induce a beneficial mutation (Charlesworth and Charlesworth 1973; Charlesworth 1974; Kirkpatrick and Barton 2006; Kirkpatrick 2010; Guerrero et al. 2012; Charlesworth and Barton 2018; Kapun and Flatt 2019; Durmaz et al. 2020; Mackintosh et al. 2022). Indeed, beginning with Dobzhansky’s seminal observations in *Drosophila pseudoobscura* (Dobzhansky 1943, 1947, 1948, 1950; Wright and Dobzhansky 1946), many inversion polymorphisms subject to spatially and/or temporally varying selection have been identified, from plants to humans (Krimbas and Powell 1992; Hoffmann et al. 2004; Stefansson et al. 2005; Hoffmann and Rieseberg 2008; Lowry and Willis 2010; Kapun et al. 2016a; Wellenreuther and Bernatchez 2018; Faria et al. 2019; Kapun and Flatt 2019; Machado et al. 2021; Lange et al. 2022).

Despite over 100 years of research, however, many fundamental questions about the adaptive role of inversions remain poorly understood (Kirkpatrick and Barton 2006; Kirkpatrick and Kern 2012; Kapun and Flatt 2019). Does adaptive divergence between inverted and standard chromosomes accumulate after an initially rare inversion has become established in a population? For instance, if an inversion has direct fitness consequences because it causes a deletion or gene expression changes near the breakpoints, we might expect that adaptive divergence between the chromosomal arrangements postdates the initial establishment of the inversion. Alternatively, do adaptive haplotypes predate the mutational origin of an inversion and then get captured by it (Kirkpatrick and Barton 2006; Kirkpatrick and Kern 2012; Guerrero et al. 2012; Charlesworth and Barton 2018; Schaal et al. 2022; Mackintosh et al. 2022)? What forms of balancing selection maintain inversion polymorphisms (Faria et al. 2019; Kapun and Flatt 2019)? And what are the genic targets of selection carried by adaptive inversions?

A promising, tractable model for tackling some of these major questions is the vinegar fly *Drosophila melanogaster*: it harbors several apparently balanced inversion polymorphisms that form parallel latitudinal clines on multiple continents (Mettler et al. 1977; Knibb et al. 1981; Knibb 1982, 1983; Lemeunier and Aulard 1992; Fabian et al. 2012; Kapun et al. 2014, 2016a, 2020; Kapun and Flatt 2019). The best studied inversion in this species is *In(3R)Payne*, a 8-Mb large paracentric inversion that spans roughly one third of the right arm of the third chromosome (*3R*; encompassing ∼1200 genes) and whose frequency varies latitudinally on several continents, most prominently along the North American and Australian east coasts (Ashburner and Lemeunier 1976; Mettler et al. 1977; Knibb et al. 1981; Knibb 1982, 1986; Lemeunier and Aulard 1992; Sezgin et al. 2004; Fabian et al. 2012; Rane et al. 2015; Kapun et al. 2014, 2016a, 2020; Kapun and Flatt 2019). The *3R Payne* inversion originated in sub-Saharan Africa >120 kya (Corbett-Detig and Hartl 2012); it thus predates the out-of-Africa expansion of *D. melanogaster* ∼4-19 kya and its subsequent colonization of other continents (Lachaise et al. 1988; David and Capy 1988; Li and Stephan 2006; Keller 2007; Kapopoulou et al. 2018, 2020; Arguello et al. 2019; Sprengelmeyer et al. 2020). Several lines of genetic and phenotypic evidence – including patterns of latitudinal clinality – suggest that this chromosomal polymorphism is adaptive (Rako et al. 2006; Kennington et al. 2006, 2007; Fabian et al. 2012; Rane et al. 2015; Kapun et al. 2014, 2016a, 2016b, Durmaz et al. 2018; Kapun and Flatt 2019; Kapun et al. 2020).

The evolutionary history of this adaptive inversion raises several interesting questions. Given its parallel clinal distribution on multiple continents, being frequent (∼40-50% or higher) in subtropical and tropical climates but rare or absent in high-latitude, temperate areas around the world (Kapun and Flatt 2019), did this inversion adapt independently – and hence convergently – to similar climatic gradients on several continents? Under such a scenario of local adaptation, the allelic content of the inversion might vary among different geographical areas (Dobzhansky 1949; Schaeffer et al. 2003; Kirkpatrick and Barton 2006). Alternatively, selection might act uniformly across a broad geographic range: if so, did the inversion capture a pre-existing adaptive haplotype in its ancestral range and then invade the rest of the world, with climatic selection favoring parallel but non-convergent spatial assortment of this polymorphism on multiple continents? With appropriate data we might be able to distinguish between these scenarios. And, given its effects on multiple fitness traits, what are likely genic targets of selection harbored by the *3R Payne* inversion?

Here we address these fundamental questions by investigating the evolutionary genomics of the *3R Payne* inversion polymorphism on four continents: in its ancestral range in Africa and in derived populations in Europe, North America, and Australia. First, we seek to elucidate the adaptive genetic basis of this inversion by combining new phased sequencing data for*3R Payne* inverted and standard karyotypes isolated from North American populations in Florida and Maine with published sequencing data from the African ancestral range as well as from Europe and Australia. We use these data to investigate patterns of phylogeography, nucleotide variability, linkage disequilibrium, karyotypic divergence, and allele sharing across populations. Second, to identify potential targets of selection spanned by the inversion, we combine *F*_ST_ outlier analyses with transcriptomic analysis of karyotypes from a derived Florida population; because *3R Payne* has been implicated in climate adaptation, we performed RNA-sequencing across two developmental temperatures.

We discuss our results in the light of theoretical predictions about expected patterns of variation and divergence of inversions (Navarro et al. 2000; Guerrero et al. 2012) and balancing selection (Zeng et al. 2021) and with regard to two commonly invoked models for adaptive inversions, Dobzhansky’s epistatic coadaptation model (Dobzhansky 1948, 1949, 1950, 1951; Charlesworth and Charlesworth 1973; Charlesworth 1974; Charlesworth and Flatt 2021) and Kirkpatrick’s and Barton’s model of ‘local adaptation’ (i.e., local selection in the face of maladaptive gene flow; Kirkpatrick and Barton 2006; Charlesworth and Barton 2018; Mackintosh et al. 2022). Under both models, a possible consequence is that the same inversion is highly locally adapted and thus contains distinct sets of adaptive alleles in different populations (Dobzhansky 1948, 1950; Prakash and Lewontin 1968, 1971; Kirkpatrick and Barton 2006),

Consistent with either model, our results suggest that *In(3R)Payne* captured adaptive alleles in the ancestral African range that predate the origin of the inversion. Yet, contrary to the above-mentioned corollary, we find relatively weak differentiation among inverted chromosomes across continents. These results indicate that the adaptive allelic content of the inversion might be ancestral and shared among populations: selection appears to have favored the spatial assortment of this ancestral polymorphism on multiple continents in a parallel fashion, resulting in qualitatively identical latitudinal clines and mediating ‘global’ (species-wide) adaptation.

## Results and Discussion

Table S1 (Supplementary Material online) gives an overview of the genomic data analyzed and indicates which data subsets were used in the different analyses presented below.

Table S2 (Supplementary Material online) provides information

## *3R Payne* is of monophyletic African origin and shows weak out-of-Africa divergence

Given that *3R Payne* is a cosmopolitan adaptive polymorphism (cf. Kapun and Flatt 2019) of sub-Saharan African evolutionary origin (Corbett-Detig and Hartl 2012), we first sought to study its phylogeography. For example, major divergence of inverted chromosomes among continents could indicate that the inversion adapted independently (i.e., convergently) to similar conditions on different continents.

Divergence-based age estimation suggests that *3R Payne* has originated ∼146,000 years ago; polymorphism-based estimation indicates a median estimate of ∼129,000 years (5% confidence limit [CL]: ∼80 kya, 95% CL: ∼196 kya; Corbett-Detig and Hartl 2012). Taking the latter estimate and assuming a generation time of ∼15 generations per year (Pool 2015), this inversion is thus at least ∼1.95 x 10^6^ generations old, i.e., roughly twice the ancestral effective population size *N_e_* (∼1.0 x 10^6^ – 1.5 x 10^6^; Kreitman 1983, Matzkin et al. 2005; Shapiro et al. 2007; Campos *et al*. 2017; Kapopoulou et al. 2018). The polymorphism is therefore probably sufficiently old for homogenizing flux between inverted and standard karyotypes to have occurred via gene conversion or double crossovers: flux rates *Φ* have been estimated to be ∼10^-4^-10^-5^ for the central regions of *D. melanogaster* inversions (Payne 1924; Chovnick 1973; Navarro et al. 1997, 2000).

The age of *3R Payne* is relevant because for sufficiently old inversions (age >> *N_e_* generations) that have captured an adaptive haplotype we might expect major peaks of divergence between inverted and standard chromosomes in the center of the inversion, which are due to the interplay of homogenizing flux and divergent selection opposing recombination (Guerrero et al. 2012; also see below). Consistent with this expectation, we have previously found major peaks of divergence in the center of *In(3R)Payne* in North American samples (Kapun et al. 2016a). In further support of a selective role, latitudinal frequency clines of *3R Payne* in Europe and North America deviate from neutral expectations (Kapun et al. 2016a, 2020), and inverted and standard karyotypes differ in their effects on several major fitness traits including body size, cold shock survival and lifespan (Rako et al. 2006; Kapun et al. 2016b; Durmaz et al. 2018).

To study the phylogeography of *3R Payne*, we investigated phylogenetic relationships among karyotypes using sequencing data from 485 strains across four continents, including data from the ancestral African range (Siavonga, Zambia; Pool et al. 2012; Lack et al. 2015, 2016; Sprengelmeyer et al. 2020) and from several derived populations in Europe (*n*=3), North America (*n*=3) and Australia (*n*=2) (fig. 1A; supplementary table S1, Supplementary Material online). Our analyses complement those of Corbett-Detig and Hartl (2012) who had examined the phylogenetic history of *In(3R)Payne* and other inversions using several African populations and single populations from Europe (France) and North America (North Carolina, USA). Based on the average number of pairwise nucleotide differences per site (nucleotide diversity *π*; Nei and Li 1979) in 100 kb non-overlapping windows, we constructed a neighbor-joining haplotype network of inverted and standard chromosomes using the Neighbor-Net method (Bryant and Moulton 2004; fig. 1B). In contrast to neighbor-joining trees, Neighbor-Net allows one to represent conflicting signals in the data, for example due to recombination.

**Fig. 1.**
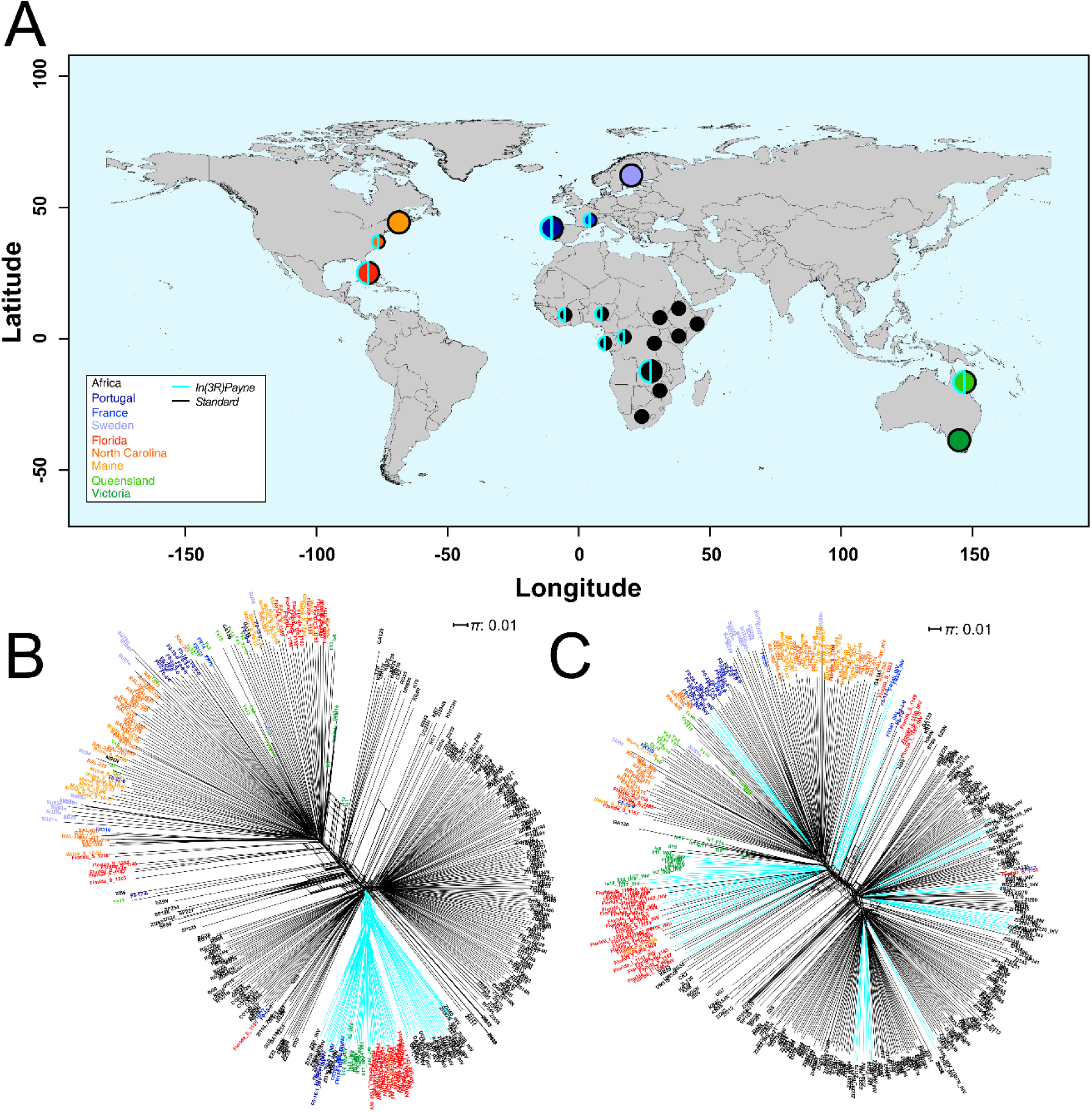
Distribution of samples and phylogenetic relationships among *In(3R)Payne* karyotypes across four continents. (A) Geographic origin of the samples used in this study. The color code indicates the continent where flies were sampled (Africa, Europe, North America, Australia). The outline of the circles indicates whether the samples contain chromosomes with *In(3R)Payne* (in cyan) and/or with the standard arrangement (in black); the size of the circles indicates whether samples were used only for phylogenetic reconstruction (small circles) or in addition also for karyotype-specific genomic analyses (large circles). (B) Haplotype network constructed from 3766 SNPs within the breakpoints of *In(3R)Payne;* cyan edges represent samples with *In(3R)Payne,* whereas black edges represent samples with the standard arrangement. (C) Haplotype network based on 4849 randomly drawn SNPs at a distance of >200 kb from *In(3L)P* and *In(3R)Payne* (see Materials and Methods). See table 1 for statistical analyses. Note that several haplotypes from Florida cluster with the NG9 reference strain (see fig. 1B, fig. 1C). This may be an artifact of our bioinformatic method for haplotype reconstruction (see Materials and Methods); we therefore excluded these samples from downstream analyses.

**Table 1.**
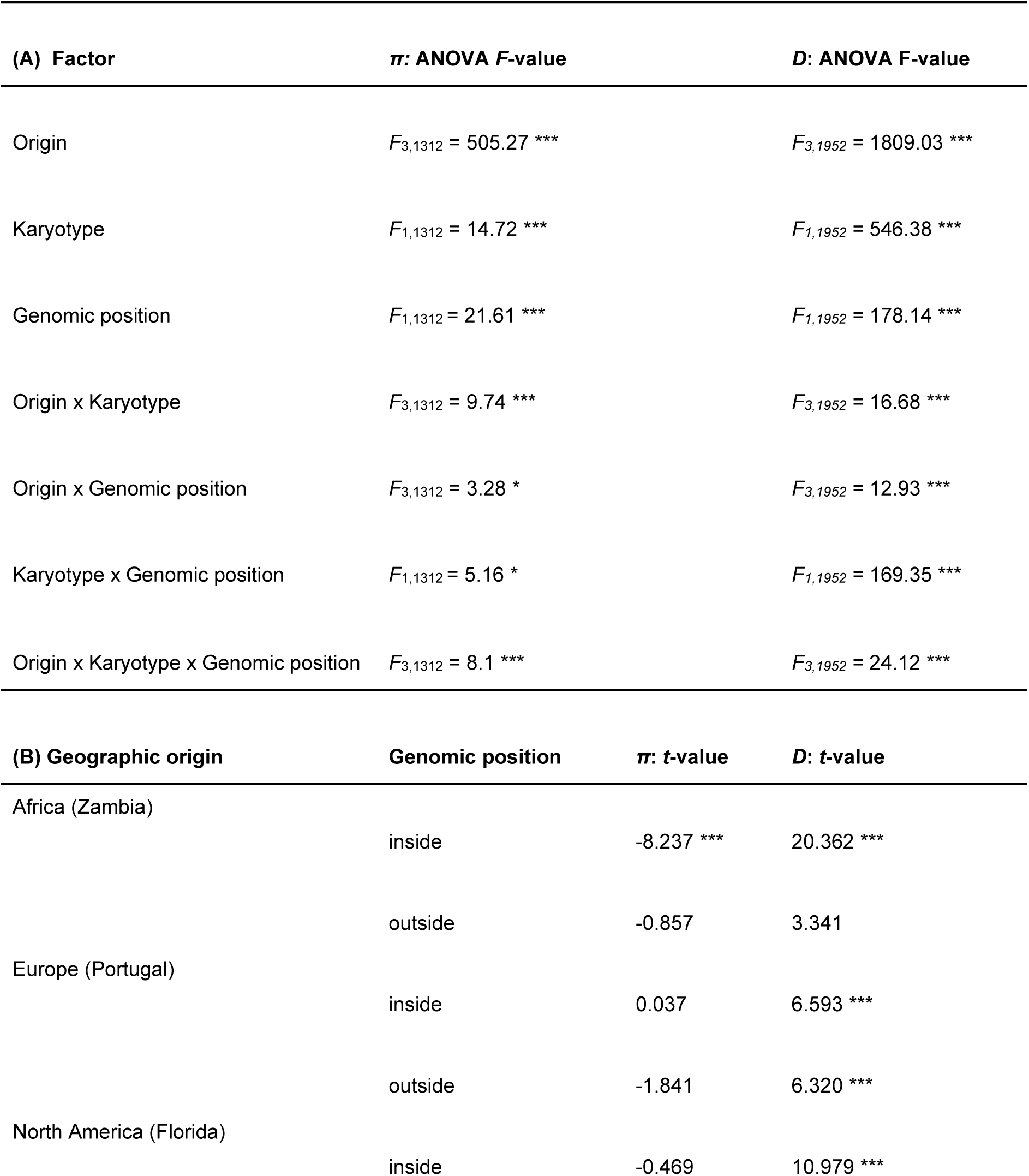
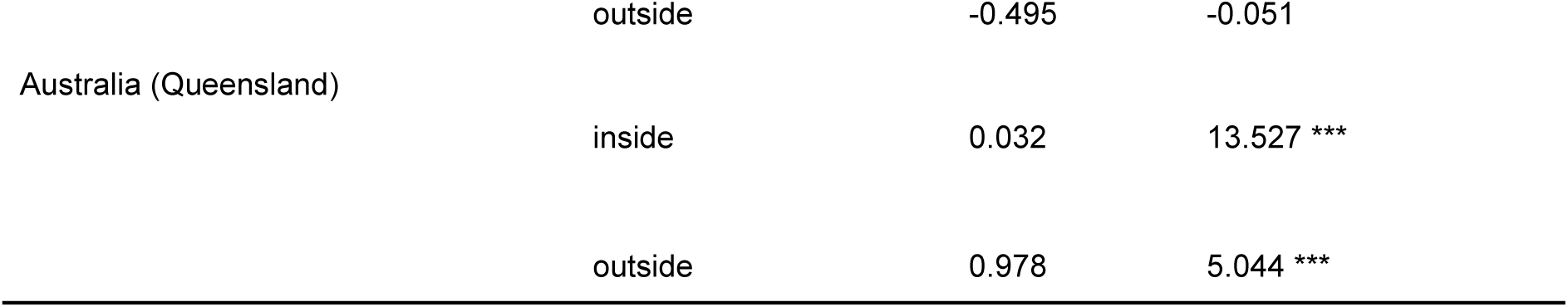
Effects on patterns of genetic variation. (A) *F*-values from a three-way ANOVA testing for differences in *π* and Tajima’s *D* with respect to geographic origin, *In(3R)Payne* karyotype, and genomic position relative to the inversion (inside vs. outside). (B) Planned contrasts based on estimated coefficients from ANOVA, testing for differences in *π* and Tajima’s *D* between inverted and standard chromosomes with respect to geography and genomic position (inside vs. outside), using the *emmeans* package in *R*. * *p* < 0.05; ** *p* < 0.01; *** *p* < 0.001. Also see fig. 1; see Materials and Methods for further details.

Inverted karyotypes clustered monophyletically within Africa, irrespective of their worldwide sampling location (fig. 1B; table 1), confirming the finding that *3R Payne* arose in sub-Saharan Africa (Corbett-Detig and Hartl 2012). This differs markedly from the pattern observed when analyzing a network based on a random set of third-chromosome SNPs at a distance > 200 kb from *In(3R)Payne* (and from the second major inversion on chromosome 3, *In(3L)P*; see Materials and Methods): here, the network structure mainly reflects geography, not *3R Payne* karyotype (fig. 1C; table 1). However, there is nonetheless a weak signal of clustering of inverted chromosome in this analysis, suggesting that the effect of *In(3R)Payne* on genetic variation might go well beyond its breakpoints (cf. Corbett-Detig and Hartl 2012).

In addition to the dominant signal of monophyletic divergence between inverted and standard karyotypes, we also found a weaker signal of geographical substructure within the inverted and standard clusters of chromosomes, indicating some divergence within karyotypes among continents (fig. 1B; table 1). The observation of substructure within the inverted karyotype bears on the question of whether *3R Payne* inverted chromosomes might be locally adapted. Under both the ‘local adaptation’ model and the epistatic coadaptation model mentioned above, loci within the inverted karyotype may be differentiated among populations if gene flow among populations disrupts locally adapted haplotypes and generates maladaptive genotypes (Prakash and Lewontin 1968, 1971; Schaeffer et al. 2003). The fact that inverted *3R Payne* chromosomes exhibit some divergence among continents is consistent with this expectation (but see results and discussion below).

### Patterns of variation are consistent with a balanced inversion polymorphism

According to coalescent models by Navarro et al. (2000) (also cf. Zeng et al. 2021), a newly arisen inversion subject to balancing selection eliminates substantial amounts of variation across a large chromosomal segment via a partial selective sweep as it increases in frequency; during the subsequent slow process of convergence to mutation-drift-flux equilibrium, variation at the breakpoints is greatly reduced as compared to the central region of the inversion where variation is restored. This is because the rate of gene flux in the form of crossing over is very low in regions close to the breakpoints and hence the effect of the partial sweep is greater. [Generally, pairing in heterokaryotypes is strongly reduced at the breakpoints, with recombination rates being very low (<<10^-4^; Navarro et al. 1997, 2000); for an inversion heterokaryotype in *D. subobscura*, Rozas and Aguadé (1994) estimated a value of 10^-7^ near the breakpoints.] By contrast, old inversions that have reached mutation-drift-flux equilibrium can exhibit greater variation at the breakpoints as compared to the inversion body (Navarro et al. 2000; cf. Wallace et al. 2013; Charlesworth 2023). This is because, over time, genetic differences between inverted and standard karyotypes become homogenized by gene flux, but this effect is much stronger in the central regions of the inversion than at the breakpoints where flux is effectively suppressed. At least 10^7^ generations are required to reach mutation-drift-flux equilibrium (Navarro et al. 2000). We thus sought to examine *π* inside and outside of the inverted region among inverted and standard *3R* chromosomes and compare our data to the expectations of Navarro et al. (2000) and Zeng et al. (2021).

Nucleotide variability on *3R* was markedly higher in the African population sample from Zambia as compared to the samples from derived population, consistent with the out-of-Africa bottleneck (Li and Stephan 2006; Lack et al. 2016; Arguello et al. 2019; Kapopoulou et al. 2020; Kapun et al. 2020, 2021) (fig. 2; supplementary fig. S1, Supplementary Material online). Inside the inverted region of African chromosomes, *π* was higher in standard relative to inverted chromosomes, but not different between arrangement types in derived populations (fig. 2; supplementary fig. S1, Supplementary Material online; see below).

**Fig. 2.**
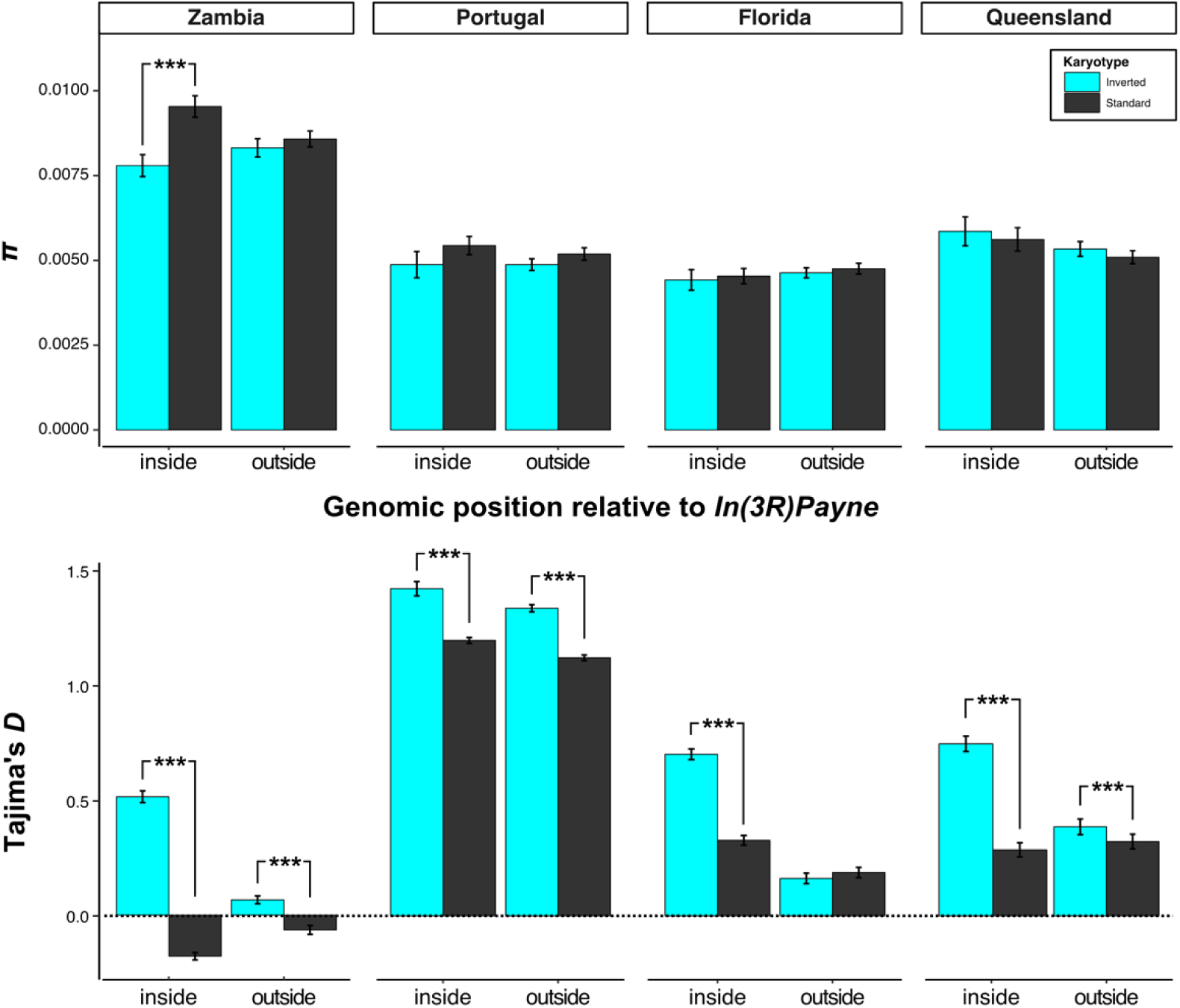
Patterns of nucleotide variability (*π*) and Tajima’s *D* in the region spanned by *In(3R)Payne.* (A) Average values of nucleotide variability *π*, calculated in 100 kb non-overlapping windows, with respect to geographic origin and genomic position relative to *In(3R)Payne,* separately for inverted and standard arrangement chromosomes. (B) Average values for Tajima’s *D*, calculated in 100 kb non-overlapping windows, separately for the two arrangement types. See table 1 for details of ANOVA results for *π* and Tajima’s *D;* asterisks (***, *p* < 0.001) represent significant *p*-values from planned contrasts. Also see supplementary fig. 1 and supplementary fig. 2 (Supplementary Material online).

In both Africa and derived populations, *π* was markedly reduced in the breakpoint regions as compared to the inversion body, resulting in a dome-shaped pattern (supplementary fig. S1, Supplementary Material online). This dome-shaped pattern agrees qualitatively well with the predictions of Navarro et al. (2000) for an inversion subject to balancing selection and which might not have reached equilibrium (i.e., inversion age < 10^7^ generations). On the other hand, consistent with equilibrium (long-term) balancing selection, *π* was higher for African standard chromosomes inside as compared to outside the inverted region (fig. 2; also see supplementary fig. S1 B, Supplementary Material online). Assuming that the frequency of the inversion is substantially lower than that of the standard arrangement, such a pattern might be expected under an equilibrium model of balancing selection with recombination (Zeng et al. 2021). Under such a scenario, the presence of the inversion would increase diversity due to the accumulation of new mutations that distinguish inverted and standard chromosomes; the coalescent time would be somewhat increased for standard alleles, due to the partial population subdivision created by the inversion, while the coalescent time would be reduced for inversion alleles (cf. Zeng et al. 2021). Recent calculations by Charlesworth (2023), which are based on our data in Fig. S1B, are consistent with *3R Payne* representing a balanced polymorphism which has reached mutation-drift-recombination equilibrium with respect to neutral or nearly neutral variants (see Navarro et al. 2000); this also suggests that *3R Payne* might be older than previously estimated (see above).

The absence of clear differences in *π* for non-African *3R* chromosomes may be due to the interplay of sufficient time for gene flux having homogenized variation between karyotypes (already in Africa), selection, and the out-of-Africa bottleneck. The fact that levels of variation in derived populations are very similar between standard and inverted chromosomes could imply that a substantial number of individuals carrying *3R Payne* has migrated out of Africa during the range expansion.

We next examined patterns of Tajima’s *D* to test for departures of the site frequency spectrum from an equilibrium standard neutral model (Tajima 1989). Relative to standard neutral expectation (*D*=0), positive values of *D* indicate an excess of intermediate frequency variants and might be consistent with a bottleneck or balancing selection; by contrast, negative values of *D* indicate an excess of rare variants which may result from a recent population expansion, from purifying selection, or from the affected genomic region having recovered variation after a selective sweep (Innan and Stephan 2000; Wallace et al. 2013; Fijarczyk and Babik 2015). Figure 2 shows average *D*, estimated separately for the two karyotypes; supplementary fig. S2 (Supplementary Material online) displays *D* along the chromosome, separately for each karyotype as well as for the pooled sample of inverted and standard chromosomes.

Average levels of *D* were positive and significantly higher for inverted as compared to standard chromosomes within populations on all continents, both inside and outside the inverted region (fig. 2; also see supplementary fig. S2, Supplementary Material online). Inverted chromosomes therefore harbor a greater frequency of intermediate variants than standard chromosomes. Positive *D* values for inverted chromosomes could arise from a bottleneck affecting the inversion within the ancestral range prior to the range expansion. But this seems unlikely as such a bottleneck should have genome-wide effects beyond the inversion; yet the positive values of *D* in the inverted region deviate markedly from the average value of *D*≍ 0 on chromosome *3R* and the genome-wide average of *D* for populations in Africa, Europe and North America (Kapun et al. 2020). Also, given that a new inversion is initially genetically invariant, one would expect more low-frequency variants as the inversion accumulates new mutations. Other possibilities might involve an incomplete sweep, or even a balanced polymorphism, among inverted chromosomes; the latter could account for the relatively high diversity of inverted chromosomes. Finally, associative overdominance (AOD), reflecting reduced recombination experienced by the inversion overall, could be involved; AOD might generate a pattern of pseudo-overdominance (Frydenberg 1963; Sved 1968; Ohta 1971; Zhao and Charlesworth 2016; Charlesworth and Charlesworth 2018; Becher et al. 2020; Gilbert et al. 2020; Berdan et al. 2021; Waller 2021; Charlesworth and Jensen 2021; Charlesworth 2022). However, under AOD, low-recombination regions still exhibit a skew towards low-frequency variants (Becher et al. 2020), so that this scenario seems improbable.

In the pooled sample of inverted and standard chromosomes (supplementary fig. S2, Supplementary Material online), we did not find evidence for positive *D* values consistent with balancing selection. Thus, our analyses of *D* are somewhat difficult to interpret; a complication with interpreting Tajima’s *D* is that it can be strongly influenced by sample size, the number of segregating sites, and by demography.

Nevertheless, several lines of evidence strongly support the notion that *In(3R)Payne* represents a balanced polymorphism, including our analyses of nucleotide variability above. For example, consistent with some form of balancing selection, *In(3R)Payne* segregates at intermediate frequencies in subtropical/tropical populations around the world: for example, the inversion attains an average frequency of ∼45% in subtropical southeastern North America and ∼60% in tropical Australian populations (Lemeunier and Aulard 1992; Rako et al. 2006; Rane et al. 2015; Kapun et al. 2016; see meta-analysis in Kapun and Flatt 2019). In Afrotropical populations, the average inversion frequency is ∼10-13%, with the highest value (∼64%) in tropical Ivory Coast (Aulard et al. 2002; Kapun and Flatt 2019). Temperate, high-latitude populations, by contrast, are fixed for the standard arrangement (Lemeunier and Aulard 1992; Kapun et al. 2016; Kapun and Flatt 2019; Kapun et al. 2020). These frequency clines, presumably in the face of sufficient gene flow to homogenize arrangement frequencies, suggest that *3R Payne* represents a balanced polymorphism driven by selection in / across heterogeneous environments (Levene 1953).

The fact that different low-latitude populations exhibit different intermediate inversion frequencies is consistent with epistatic coadaptation: under such a model, there exist multiple equilibria and quasi-equilibria for the frequency of the inversion, and the frequency which it will ultimately attain will depend on the history, the initial conditions of the population, and/or the local environment (Charlesworth 1974; also see Dobzhansky and Pavlovsky 1957; Lewontin 1974). Although the model of Charlesworth (1974) assumes constant fitness values, it leads to apparent frequency-dependent selection. Interestingly, Nassar et al. (1973) found that negative frequency-dependent viability selection operates on *In(3R)Payne* under crowded larval conditions, giving further credence to a scenario of balancing selection.

Some studies have reported that *In(3R)Payne* can locally reach near-fixation or fixation in some Australian populations (Knibb et al. 1981; Anderson et al. 2005; Umina et al. 2005), an observation that seems to be at odds with a balanced polymorphism. However, the sample size in the study of Knibb et al. (1981) was extremely low. Moreover, drift can cause the fixation of one variant and loss of the alternative variant despite balancing selection (Robertson 1962; Ewens and Thomson 1970). Also, the selective factors favoring the polymorphism might be environmentally sensitive, so that balancing selection could break down in some locations.

Overall, the data available to date indicate that *In(3R)Payne* segregates at intermediate frequencies in the majority of low-latitude populations around the world and that fixation of the inversion is rare (Kapun and Flatt 2019) – this pattern and our above results are thus broadly consistent with balancing selection and/or spatially varying selection (Levene 1953) maintaining this polymorphism.

### Patterns of LD are compatible with linked selection maintaining the inversion

Next, we examined patterns of linkage disequilibrium (LD). Three aspects of LD can be distinguished: (i) LD among markers without reference to karyotype; (ii) LD between a marker and inverted vs. standard arrangements; and (iii) LD between markers within inverted or within standard chromosomes. Because inversions strongly reduce the products of recombination in heterozygous state, heterokaryotypes (or pools of inverted and standard chromosomes) should exhibit increased LD as compared to homokaryotypes (aspect ii); for sufficiently old inversions that evolve neutrally we might expect that LD decreases towards the center of the inversion due to gene flux between the karyotypes, even though such a pattern is difficult to distinguish from direct positive selection at the breakpoints (Navarro et al. 1997; Guerrero et al. 2012). Within the class of inverted chromosomes (aspect iii), LD can be higher than within standard chromosomes because of a smaller *N_e_* of the inversion.

LD between the *3R Payne* inversion and marker loci has been previously studied by Kojima et al. (1970), Langley et al. (1974), Voelker et al. (1978), Sezgin et al. (2004) and Kennington et al. (2006); such LD between the inversion and marker loci might be due to hitchhiking of a neutral variant initially associated with the inversion by chance (Ishii and Charlesworth 1977) or due to subsequent new mutations that differentiate the karyotypes. More recently, Rane et al. (2015) examined LD associated with *In(3R)Payne* in Australian samples using RAD-sequencing data. Here, we sought to use phased genomic data to compare patterns of LD in the region spanned by *3R Payne* in African, European, North American and Australian samples with single nucleotide resolution (fig. 3; supplementary fig. S3, Supplementary Material online).

**Fig. 3.**
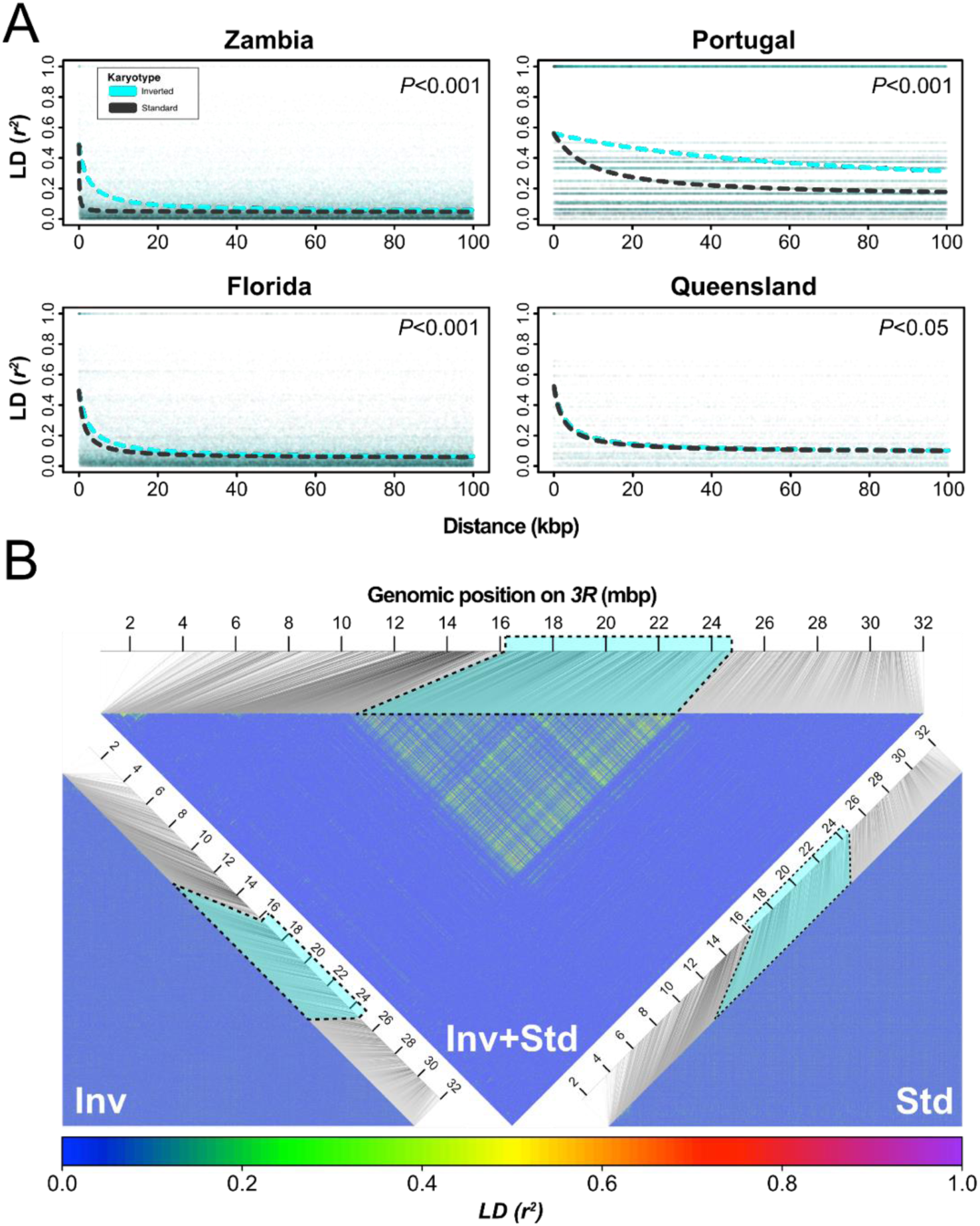
Patterns of short-and long-range linkage disequilibrium (LD) in the region spanned by *In(3R)Payne.* (A) Distribution and decay of LD as estimated by *r*^2^ within 100 kb distance, based on 5000 randomly drawn SNPs inside the region spanned by *In(3R)Payne* for standard (grey) and inverted (cyan) chromosomes from different geographic samples. Significant *p*-values in the top right corners of the plots indicate differences in the decay of LD among karyotypes as inferred from analyses of deviance applied to non-linear regression models (see Material and Methods for details). (B) Triangular heat maps with estimates of *r*^2^ for 5000 SNPs randomly drawn from chromosomal arm *3R* in samples from North America (Florida). We restricted our analyses to subsamples of 5000 SNPs due to computational reasons: *n* = 5000 SNPs implies *n*(*n*-1)/2 = 12,497,500 pairwise comparisons, with much larger numbers becoming computationally prohibitive. In the upper triangle, *r*^2^ was estimated jointly for inverted and standard chromosomes (see fig. S3, Supplementary Material online, for similar plots for the other continents). The bottom left and right plots show separate *r*^2^ estimates for inverted and standard chromosomes, respectively.

As compared to standard chromosomes, inverted arrangements showed significantly higher levels of short-range LD (*r*^2^) within the region spanned by *In(3R)Payne* and a slower decay of LD with physical distance (fig. 3A). A plausible explanation is that this pattern is due to drift within the two semi-isolated subpopulations of inverted vs. standard chromosomes, with inverted chromosomes exhibiting both lower recombination and lower *N_e_*. The pattern of decay was similar for inverted chromosomes across continents, except for the Portuguese sample, perhaps due to the rather small number of sampled chromosomes and the overall lower frequency of *3R Payne* in Europe (Kapun et al. 2020). By contrast, the pattern of decay for standard chromosomes differed markedly among continents: while in the African sample LD leveled off to *r*^2^ < 0.1 within a few hundred base pairs (bp), the decay of LD in standard arrangements from North America (Florida) and Australia (Queensland) closely resembled that of inverted chromosomes (fig. 3A), probably reflecting bottlenecks in the derived samples. The patterns for the derived populations were generally less clear than for the Zambian population, presumably due to the out-of-Africa bottleneck.

Next, we examined long-range LD (fig. 3B; supplementary fig. S3, Supplementary Material online). We first analyzed LD within each karyotype. For both standard and inverted arrangements, LD levels did not exceed *r*^2^ > 0.1 within distances of a few kbp, revealing long-range linkage equilibrium within karyotypes. In marked contrast, when jointly analyzing the pool of inverted and standard karyotypes from Florida (Fig. 3B), we observed strong long-range LD within the inverted region, involving SNPs several million bp away from each other and suggesting that major associations among loci are driven by hetero-not homokaryotypes. These patterns were similar for the other continents, with major LD between but not within karyotypes (supplementary fig. S3, Supplementary Material online). Likewise, no strong LD was seen within Australian karyotypes; this is contrary to Rane et al. (2015) and likely due to a misclassification of karyotypes in that study (see below; Materials and Methods; supplementary fig. S4, Supplementary Material online).

Notably, in European, North American and Australian samples we found large clusters of SNPs in the center of the inversion that are in strong LD with each other and the proximal and distal breakpoints, interspersed by regions of low or no LD (fig. 3B; supplementary fig. S3, Supplementary Material online). For Australia, our data agree well with those of Kennington et al. (1997) who found LD among marker loci within and near *In(3R)Payne* and between these loci and the inversion itself, including marked associations in the center of the inversion. In the African sample, we also observed LD between the breakpoints and center regions of elevated LD, yet these central clusters of high LD were much less prominent than in the derived populations (supplementary fig. S3, Supplementary Material online). These patterns of long-range LD almost certainly reflect the strong divergence between inverted and standard arrangements (cf. Zeng et al. 2021); the clearer patterns seen for non-African populations might be due to lower diversity which tends to sharpen up divergence patterns (Nordborg et al. 1996).

Associations between an inversion and loci within the inverted region can have several causes that are difficult to distinguish (Strobeck 1983; Navarro et al. 1996): the inversion might have become associated with neutral alleles when it formed (Ishii and Charlesworth 1977; Nei and Li 1980), or it might be linked to neutral loci subject to drift (Nei and Li 1975; Strobeck 1983); or selection might maintain associations between selected loci spanned by the inversion and the inversion itself despite flux between arrangements (see above; also see Prakash and Lewontin 1968; Charlesworth and Charlesworth 1973; Charlesworth 1974; Ishii and Charlesworth 1977; Schaeffer et al. 2003; Guerrero et al. 2012; Fuller et al. 2017). The extent of such associations depends on the flux rate, the effective number of inverted and standard chromosomes, and the inversion age (Ishii and Charlesworth 1977; Nei and Li 1980). Theory suggests that the half-life of decay of an association between a neutral locus and an inversion is on the order of the reciprocal of the flux rate in heterokaryotypes (Ishii and Charlesworth 1977; Nei and Li 1980). Selection can retard this decay considerably, but only when the neutral locus is very closely linked to one of the selected loci involved in maintaining the polymorphism (Ishii and Charlesworth 1977).

How do our data compare to these predictions? Assuming a gene flux rate *Φ* of ∼10^-5^ in the center of the inversion (Chovnick 1973), the timescale for the decay of the association would be on the order of ∼10^5^ generations (∼7000-10,000 years, assuming 10-15 generations per year). Given that *In(3R)Payne* is at least ∼129k years old and globally quite frequent (Kapun and Flatt 2019: average global frequency ∼15%, based on 530 samples from 34 independent studies spanning >50 years of data), and given that *N_e_* is large (∼10^6^), gene flux should have had ample opportunity to break down strong LD associated with this inversion. Our data are thus consistent with the selective maintenance of the center peaks inside the inversion. On the other hand, *D. melanogaster* has undergone an expansion from southern-central Africa and a major out-of-Africa bottleneck; began to spread from the Middle East into Europe ∼1800 years ago; and colonized North America and Australia ∼100-150 years ago (Hoffmann and Weeks 2007; Keller 2007; Sprengelmeyer et al. 2020): demographic events such as bottlenecks, drift and/or founder effects can therefore not be ruled out as having influenced LD associated with *In(3R)Payne*.

### Major peaks of divergence inside the inversion are shared across continents

To study chromosome-wide patterns of differentiation as a function of *3R Payne* karyotype we used *F*_ST_, a normalized measure of pairwise allele frequency differentiation (Weir and Cockerham 1984). [In the context of karyotypic differentiation, it would be more accurate to call this quantity *F*_AT_, a measure of variation between allelic classes at a polymorphic locus; Charlesworth et al. 1997.] We were particularly interested in determining whether there might be major peaks of divergence between standard and inverted chromosomes in the center of the inverted region, away from the breakpoints. For sufficiently old inversions, and assuming the existence of targets of selection within the inversion, coalescent theory predicts that selection might lead to peaks inside the inversion body, which are centered on the adaptive loci and selectively maintained in the face of homogenizing flux between standard and inverted chromosomes (Guerrero et al. 2012). This pattern is not unique to inversions: any form of balancing selection will lead to a peak of divergence and LD around the target of selection at equilibrium (Hudson and Kaplan 1988; Kaplan et al. 1988; Guerrero et al. 2012; Zeng et al. 2021). For old inversions, this leads to a characteristic pattern of divergence between the karyotypes (Guerrero et al. 2012; Kirkpatrick 2017): the pattern of divergence resembles the cables of a suspension bridge with peaks of divergence both at the breakpoints (where recombination is greatly reduced) and in the center of the inversion (where selection opposes recombination). Such center peaks of divergence could arise from either the Kirkpatrick-Barton model or the epistatic coadaptation mechanism (Guerrero et al. 2012; Charlesworth and Barton 2018; Kapun and Flatt 2019; Durmaz et al. 2020; Charlesworth and Flatt 2021); sweeps within inverted or standard chromosomes could also generate such peaks. We have previously found such peaks in pool-sequencing data for North American samples (Kapun et al. 2016a), and Rane et al. (2015) had examined such peaks in Australian data using RAD-sequencing.

Here, we sought to revisit these results and to extend them to African and European samples. Secondly, we aimed to assess the contribution of *3R Payne* to divergence across latitudinal clines in Europe, North America and Australia (Kolaczkowski et al. 2011; Fabian et al. 2012; Rane et al. 2015; Kapun et al. 2016a, 2020). To this end, we studied the effects on divergence of ‘karyotype’ (‘K’, comparing inverted vs. standard arrangements within the same low-latitude populations), ‘geography’ (‘G’, comparing standard chromosomes between low-and high-latitude populations) and ‘geography plus karyotype’ (‘G+K’, comparing low-latitude inverted chromosomes with high-latitude standard chromosomes) (see Materials and Methods). Figure 4 shows patterns of *F*_ST_ for these effects as a function of position on *3R*, including estimates of LD between SNPs in the region spanned by the inversion and the inversion itself. Inspection of these patterns revealed several interesting findings.

**Fig. 4.**
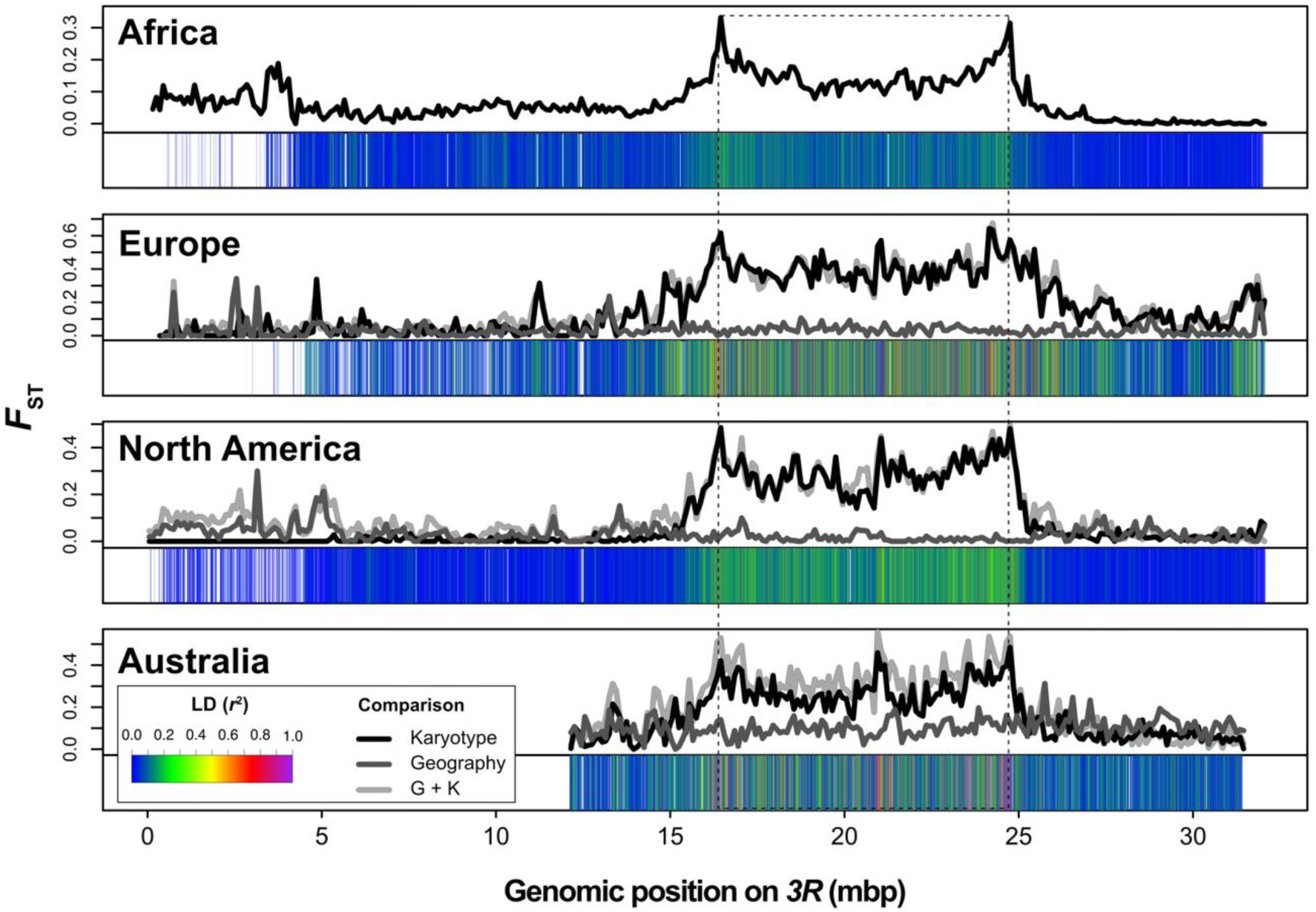
Chromosome-wide patterns of genetic differentiation (*F*_ST_) due to *In(3R)Payne* karyotype and/or the effects of geography. Line plots show the distribution of *F*_ST_ in 100 kb non-overlapping windows along chromosome arm *3R*. For the non-clinal African population sample from Zambia (top panel), the line plot shows *F*_ST_ between inverted and standard chromosomes within the Siavonga population sample. For the non-African populations from Europe, North America and Australia, which are all situated along latitudinal gradients, the different lines depict the effects of ‘karyotype’ (black line; pairwise differences between standard and inverted chromosome from within a given low-latitude population), ‘geography’ (dark grey line; pairwise comparisons between standard arrangement chromosomes from low– vs. high-latitude populations, i.e. from the cline ‘ends’); and of ‘geography plus karyotype’ (‘G+K’; light grey line; pairwise comparisons of inverted and standard chromosomes between the endpoints of a given cline). Heat maps beneath each line plot show *r*^2^ between each SNP and *In(3R)Payne.* Note that genomic information for the first 12 million bp is not available for the Australian data.

First, we observed marked divergence in the region spanned by the inversion between inverted and standard chromosomes on all four continents (effect of ‘K’), with pronounced peaks in the breakpoint regions (fig. 4, black lines). For derived populations, where *3R Payne* exhibits latitudinal clines on different continents, this divergence is similar when contrasting inverted and standard chromosomes from within the same low-latitude populations (effect of ‘K’) and when comparing low-latitude inverted with high-latitude standard arrangements (effect of ‘G+K’, comparing karyotypes between the cline ‘ends’; fig. 4, light grey lines). By contrast, divergence is low when comparing standard chromosomes between low-and high-latitude populations in Europe, North America and Australia (effect of ‘G’; fig. 4, dark grey lines). This is also quantified for derived populations in fig. 5 and table 2, for both the region inside and outside of *In(3R)Payne*.

**Table 2.** Effects of *In(3R)Payne* karyotype and/or geographic origin on pairwise *F*_ST_ differences. *F*-values from a one-way ANOVAs testing for differences in divergence using pairwise *F*_ST_ comparisons as input see Materials and Methods for details). ANOVAs were performed separately for each continent and genomic region (inside vs. outside inversion) with respect to *In(3R)Payne.* To determine which of the three levels (Karyotype; Geography; Karyotype + Geography) differ from each other we performed Tukey’s HSD post-hoc tests. * *p* < 0.05; ** *p* < 0.01; *** *p* < 0.001. Also see fig. 5; see Materials and Methods for further details.

|  |  |  | Tukey's HSD statistic |  |  |
| --- | --- | --- | --- | --- | --- |
| Origin | Position | ANOVA $F$-value | Karyotype vs. Geography | Karyotype + Geography vs. Geography | Karyotype + Geography vs. Karyotype |
| Europe | Inside | $F_{2,366} = 543.85$ *** | 0.33*** | 0.34*** | 0.01 |
| | Outside | $F_{2,780} = 8.4722$ ** | 0.01 | 0.02*** | 0.01 |
| North America | Inside | $F_{2,366} = 221.02$ *** | 0.21*** | 0.23*** | 0.02 |
| | Outside | $F_{2,792} = 119.6$ *** | -0.03*** | 0.02*** | 0.05*** |
| Australia | Inside | $F_{2,366} = 170.35$ *** | 0.14*** | 0.2*** | 0.06*** |
| | Outside | $F_{2,255} = 77.524$ *** | -0.07*** | -0.02*** | 0.05*** |

**Fig. 5.**
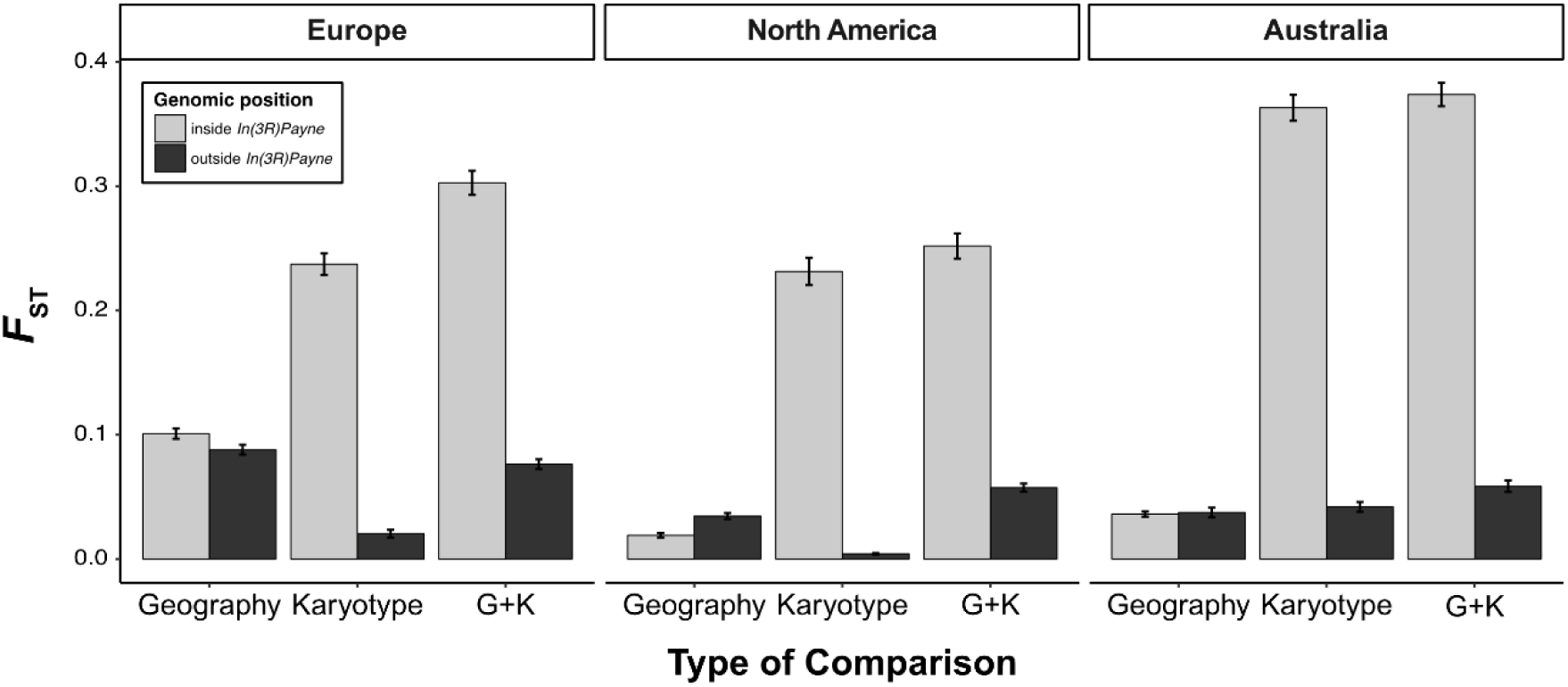
Genetic differentiation (*F*_ST_) as a function of *In(3R)Payne* karyotype and/or geography. Bar plots show average values of *F*_ST_ in 100 kb non-overlapping windows in different genomic regions relative to *In(3R)Payne* (inside vs. outside the inverted region) for the three non-African continents (Europe, North America, Australia) which include clinal (low– vs. high-latitude) populations samples. The different bars represent pairwise *F*_ST_ comparisons for (i) geographic differentiation (‘G’, comparing standard arrangement chromosomes from populations at the endpoints of clines), (ii) karyotypic differentiation (‘K’, comparing inverted and standard arrangement chromosomes sampled from within the same low-latitude populations), and (iii) geographic plus karyotypic differentiation (‘G +K’, comparing inverted chromosomes from low-latitude populations with standard chromosomes from high-latitude populations). See Materials and Methods for details; also see fig. 4 and table 2 for statistical analyses.

These results indicate that *3R Payne* karyotype is the major determinant of divergence on chromosome arm *3R* in all populations examined, and that clinal divergence in non-African populations is predominantly caused by the divergence between inversion karyotypes, not by geography; geographic differentiation inside the inverted region is much weaker than karyotypic differentiation, despite very large geographical distances (∼2600-3900 km) between the ‘endpoints’ of the clines on different continents (Kapun et al. 2016a). By contrast, outside the inverted region, patterns of divergence are consistent with isolation by distance. These results agree well with previous pool-sequencing analyses of *In(3R)Payne* in North America and Europe (Kapun et al. 2016a, 2020). However, for Australia our findings differ from those of Rane et al. (2015) who found no major effect of karyotype on divergence in the Queensland low-latitude population sample. Our reclassification of karyotypes in this dataset suggests that this previously reported pattern was due to a partial misassignment of karyotypes. Using our new classification, we found major karyotypic divergence in the Queensland sample (fig. 4; fig. 5; supplementary fig. S4, Supplementary Material online; table 2), which is fully consistent with our analyses of European and North American karyotypes and our analyses in fig. 1.

Second, coarse-grained patterns of karyotypic divergence and LD, especially for derived populations, are highly congruent across continents, including Australia (fig. 4; fig. 5; supplementary fig. S4, Supplementary Material online). The parallel divergence due to *In(3R)Payne* is underscored by strong correlations between levels of *F*_ST_ with respect to karyotype across continents, including Africa (supplementary fig. S5, Supplementary Material online). This might reflect that most SNPs are neutral and in LD with the inversion; on the other hand, it is also consistent with parallel clinal adaptation to similar environmental gradients around the world. Together with our phylogenetic analysis, this speaks against a scenario of ‘strict’ local adaptation whereby the same inversion is genetically differently adapted to distinct local conditions – under such a scenario one might expect larger geographical differentiation among inverted chromosomes (Dobzhansky 1951; Prakash and Lewontin 1968, 1971; Schaeffer et al. 2003; Kirkpatrick and Barton 2006; see below and supplementary fig. S6, Supplementary Material online).

Third, on a fine-grained scale, we observed major peaks of divergence in the inversion center that are shared among all non-African populations (fig. 4; supplementary fig. S4, Supplementary Material online). Most prominently, there is a massive central peak of divergence of ∼200-300 kb length (position on *3R*: ∼20.9 – 21.2 Mbp) that is common to derived populations in Europe, North America and Australia, with alleles in this peak being in strong LD with the breakpoints (fig. 4; also see fig. 3). These shared peaks, as well as the consistency of LD structure among populations on several continents, are consistent with the idea that linked selection maintains non-random associations between the center peaks and the breakpoints despite homogenizing flux between arrangements (see above; Charlesworth 1974; Guerrero et al. 2012; cf. Prakash and Lewontin 1968, 1971; Lewontin 1974). However, the history of these derived populations is not independent, and bottleneck events (or a strong selective sweep within inverted chromosomes) cannot be ruled out as alternative explanations.

Fourth, although these major center peaks seem to be absent in the African sample (top panel in fig. 4), preliminary results by Brian Charlesworth (personal communication) suggest that the observed *F*_ST_ between karyotypes of ∼0.1 in the African sample for sites away from the breakpoints agrees qualitatively well with expected neutral divergence between karyotypes (*F*_ST_ = 0.13), assuming an equilibrium balanced polymorphism under an island model of population structure (subdivision with neutral *F*_ST_ = 0.05; inversion frequency = 0.1; rate of exchange = 10^-6^ per site per generation). The pattern in the Zambian sample might thus be compatible with *3R Payne* representing a long-term balanced polymorphism (also see discussion of *π* above; see discussion in Charlesworth 2023). This prompted us to take a closer look at inversion-associated alleles within the ancestral African sample.

### The inversion appears to have captured adaptive alleles in its ancestral range

Several models of adaptive inversion evolution posit that a new inversion might capture a pre-existing adaptive haplotype, i.e., a set of selected loci that are in loose LD (Dobzhansky 1949, 1950, 1951; Charlesworth and Charlesworth 1973; Charlesworth 1974; Kirkpatrick and Barton 2006; Charlesworth and Barton 2018; Charlesworth and Flatt 2021; Schaal et al. 2022; also cf. Kimura 1956); alternatively, adaptive divergence between inverted and standard arrangements might have accumulated after the inversion was established. In the former case, we might expect that standard chromosomes in the ancestral African range still carry some of the presumably adaptive, pre-existing alleles that were captured by the inversion when it first arose (Kirkpatrick 2017).

Consistent with differentiation among karyotypes not being the result of (continent-specific) local adaptation but having arisen prior to the out-of-Africa migration, we failed to observe elevated divergence within the inversion body among inverted chromosomes from different continents (supplementary fig. S6, Supplementary Material online).

To further explore this idea, we quantified the frequency of inversion-specific alleles, defined as SNPs with *F*_ST_ ≥ 0.9 between inverted and standard chromosomes in the North American sample from Florida, among African (and for comparison also among European) standard and inverted chromosomes (fig. 6).

**Fig. 6.**
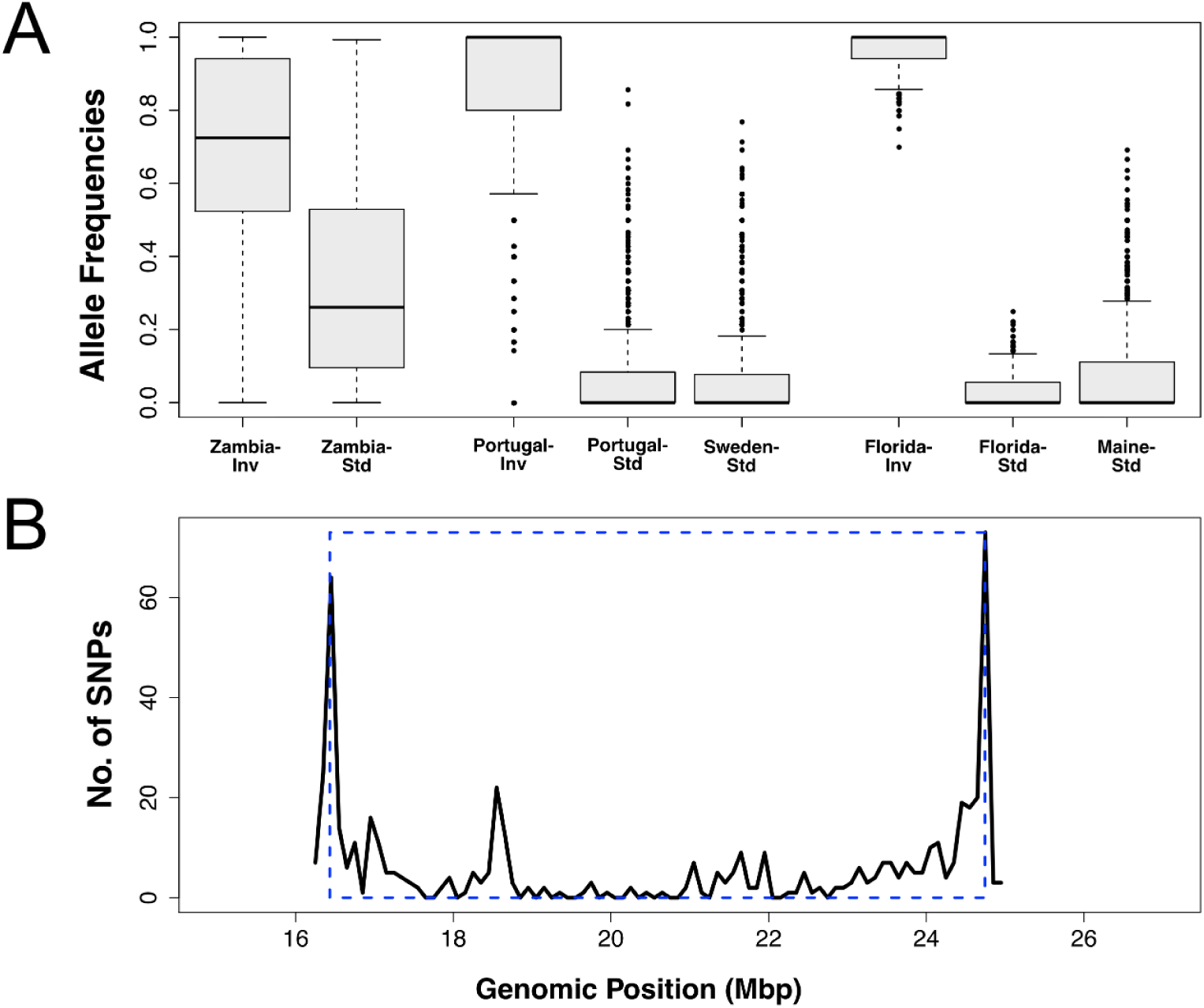
African origin of inversion-specific alleles. Panel A shows median allele frequencies of inversion-specific alleles from North America (*F*_ST_ ≥ 0.9) in inverted and standard arrangement chromosomes in population samples from Africa, Europe and North America. Panel B shows that highly differentiated SNPs in the Zambian population (exhibiting a frequency difference ≥ 0.5 between standard and inverted chromosomes) are mostly clustered around the inversion breakpoints, with some smaller clusters (‘mini-peaks’) of SNPs also visible around positions ∼19 Mbp and ∼21 Mbp. Also see supplementary fig. S7 (Supplementary Material online). The analyses above are based on 1786 SNPs in total, and 277 SNPs in the Zambia sample.

This analysis revealed that alleles that are highly ‘inversion-specific’ outside of Africa are polymorphic among *both* African standard *and* African inverted chromosomes (fig. 6A), possibly indicating that they represent ancestral alleles that have been captured by the inversion. The major enrichment of ‘inversion-specific’ alleles among African inverted relative to standard chromosomes (frequency difference of ∼45% between inverted and standard karyotypes; fig. 6A) might also be consistent with the inversion having captured these alleles before the out-of-Africa expansion. If so, this would speak against a scenario whereby the inversion spread to some appreciable frequency by drift and then gained adaptive variants via influx from the subpopulation of standard chromosomes by recombination or through new mutations, with the inversion driven to high frequency by hitchhiking.

Repeating the analysis in fig. 6A by using highly inversion-specific alleles (*F*_ST_ ≥ 0.9) as defined based on the Zambian population (instead of defining them, as above, based on the Florida sample) also revealed major frequency differentiation between African inverted and standard alleles in derived populations, consistent with the notion that African alleles underpin the divergence of *In(3R)Payne* karyotypes in derived populations (supplementary fig. S7, Supplementary Material online).

Figure 6B shows the distribution of ‘inversion-specific’ SNPs (as defined using the Florida sample) in the African sample with a frequency difference ≥ 50% between standard and inverted chromosomes: the resulting pattern delineates the breakpoints clearly, indicating that divergence in the African sample is driven by suppressed recombination at the breakpoints. Also note the two ‘mini-peak’ regions away from the breakpoints (a larger one at ∼19 Mbp; and a smaller range of peaks at ∼21 Mbp) where flux is expected to be much higher than at the breakpoints: the locations of these mini-peaks correspond well with those of the major central peaks seen in European, North American and (for the second peak region) Australian samples (fig. 4). Because levels of diversity are very similar between standard and inverted chromosomes in the derived populations, it seems improbable that these peaks are due to very low *N_e_* of inverted chromosomes leaving Africa. Nonetheless, we cannot rule out that these peaks might have become more pronounced during the range expansion, potentially due to the out-of-Africa bottleneck and/or drift, perhaps in addition to selection.

### Genetic divergence between inversion karyotypes is shared among continents

Because patterns of karyotypic divergence and LD looked very similar across continents (fig. 4), especially for derived populations, we were interested in quantifying the geographical overlap in the number of inversion-associated candidate genes and SNPs (fig. 7; candidates defined by SNPs with *F*_ST_ ≥ 0.9 between inverted and standard karyotypes; see Materials and Methods). Overall, we observed significant sharing of candidate genes and SNPs across continents (fig. 7). However, the inclusion of the Australian data resulted in very low levels of sharing, perhaps because this dataset is based on reduced-representation RAD-sequencing with low resolution; we therefore excluded the Australian data from the analysis (fig. 7). Independent of whether the Australian data were excluded or not (not shown), we identified major overlap of candidates between Europe and North America (fig. 7), probably because of the demographic and genetic similarity of populations on these continents. Importantly, when excluding the Australian data, we found a highly significant overlap of 174 candidate genes and 34 SNPs that are shared between Africa, Europe and North America (fig. 7A, fig. 7B) – these loci might thus underlie the shared pattern of karyotypic divergence across continents (supplementary table S3; supplementary table S4; Supplementary Material online; see below).

**Fig. 7.**
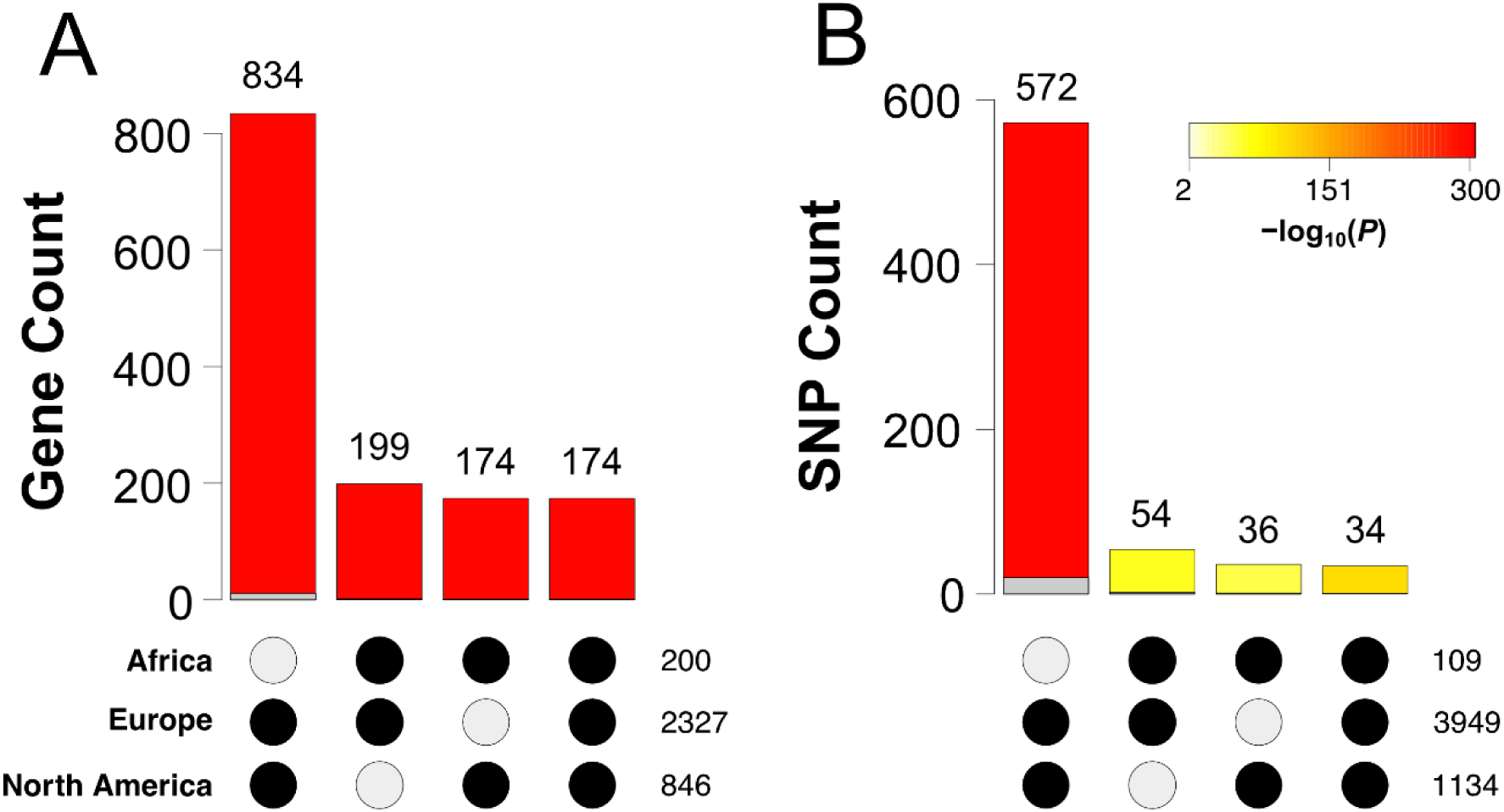
Overlap of *In(3R)Payne*-associated candidate genes and SNPs among continents. Bar plots show the counts of overlapping candidate genes (A) or SNPs (B) in Africa, North America and Europe, as indicated by black dots underneath each bar plot. The total number of candidates for each dataset is shown on the right side of the black dots. The color of the bars corresponds to the significance of the overlaps, as inferred by the *R* package *SuperExactTest* and as indicated by the color gradient in the legend. The grey overlays at the bottom of the bars indicate the amount of expected overlap. Also see supplementary table S3 and supplementary table S4 (Supplementary Material online); also see Materials and Methods.

When examining the putative functional effects of these candidate SNPs, we found a significant deficiency of SNPs in the intergenic region of African, European and North American samples and a significant enrichment in the 2 kb upstream region of genes in European and North American samples (see supplementary fig. S8; supplementary table S4; Supplementary Material online). These findings suggest that several candidate SNPs might influence gene regulation and gene expression patterns.

### The inversion affects gene expression in a temperature-dependent manner

A handful of studies has examined clinal differences in gene expression in Australian and North American *D. melanogaster* (Levine et al. 2011; Chen et al. 2012; Zhao et al. 2015; Juneja et al. 2016; Clemson et al. 2018), but whether *3R Payne* contributes to these patterns remains unknown. To investigate the effects of this inversion polymorphism on differential expression (DE), and to complement our genomic analyses, we analyzed RNA-Seq data from adult female whole-body transcriptomes of different *In(3R)Payne* karyotypes. Our dataset consisted of 9 biological replicates each for Florida inverted (FI), Florida standard (FS) and Maine standard (MS) homokaryotypes (isochromosomal lines), with each group reared at two developmental temperatures (18°C, 25°C) prior to RNA extraction (3 karyotypes x 2 temperatures x 9 replicates = 54 samples in total). Because the *3R Payne* inversion is involved in climate adaptation (Kapun et al. 2016a; Kapun and Flatt 2019), this design allowed us to examine whether developmental temperature interacts with karyotype and/or geographic origin in affecting expression (supplementary table S5, Supplementary Material online; see Materials and Methods).

Approximately 60% of all analyzed genes genome-wide (n=9724) showed significant DE in response to temperature (n=5841; Benjamini-Hochberg [BH]-corrected *p* < 0.05; supplementary table S5, Supplementary Material online), in agreement with previous work reporting high levels of expression plasticity across different rearing temperatures (Chen et al. 2015). Inversion karyotype had a much weaker effect: only 0.49% of all genes were differentially expressed between karyotypes (Florida inverted vs. standard; n=46; BH-corrected *p* < 0.05) and 0.45% in response to the effect of karyotype plus geography (Florida inverted vs. Maine standard; n=44; BH-corrected *p* < 0.05) (supplementary table S5, Supplementary Material online).

Interestingly, we failed to identify any DE in response to the effects of geography alone (Florida standard vs. Maine standard; BH-corrected *p* > 0.05; see supplementary table S5, Supplementary Material online); the effects of karyotype plus geography thus seem to be driven by karyotypic differences, not geography. We were thus interested in comparing our transcriptomic candidates to the RNA-seq data of Zhao et al. (2015) who had examined DE between populations from Panama (low latitude) and Maine (high latitude) at two growth temperatures (21°C, 29°C). This analysis revealed significant overlaps between the effects of karyotype plus geography in our data and differentially expressed genes (DEG) identified by Zhao et al. (2015) (*SuperExactTest*; *p* < 0.05; supplementary table S6; also see supplementary table S3, Supplementary Material online); overlaps between DEG found by Zhao et al. (2015) and the effects of karyotype in our data were marginally non-significant. These results, together with the analyses of Zhao et al. (2015), suggest that *In(3R)Payne* makes a major contribution to latitudinal differentiation in gene expression patterns.

We also found pervasive interactions between inversion karyotype and growth temperature: temperature had a major influence on both the magnitude of DE and the number of DEG between karyotypes (supplementary table S5, Supplementary Material online). While 648 genes were differentially expressed between inverted and standard arrangement females that had developed at 18°C, we did not find any gene exhibiting significant DE between karyotypes at 25°C (fig. 8A; supplementary table S5, Supplementary Material online). This suggests that variants associated with the inverted arrangement might be more sensitive to lower temperatures, maybe due to a loss of buffering or because of increased compensatory plasticity at 18°C (cf. Huang et al. 2022), lending further support to the role of *3R Payne* in climate adaptation (also see Pool et al. 2016).

**Fig. 8.**
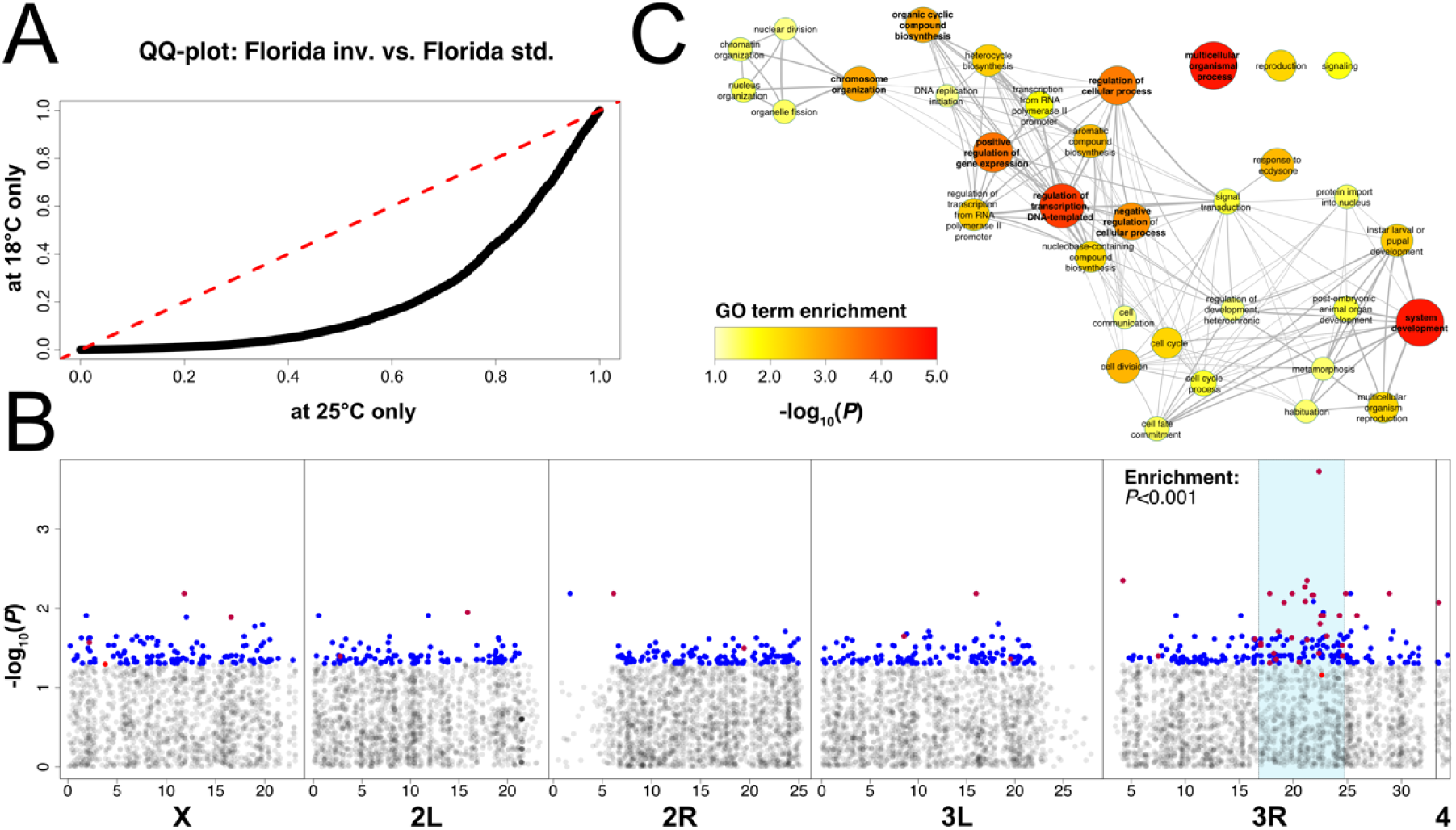
Transcriptomic analyses of *In(3R)Payne* karyotypes. (A) Q-Q plot comparing *p*-values from differential expression (DE) analyses with *limma voom* between North American *3R Payne* inverted and standard chromosomes at 18°C and 25°C, respectively. Since karyotype-specific DE was much stronger at 18°C, we focused on this dataset for downstream analyses. (B) Manhattan plot depicting – log_10_(*p*)-values for each gene relative to its average genomic position. Candidate genes with karyotype-specific expression irrespective of temperature are highlighted in red; those showing karyotype-specific expression at 18°C only in blue; and those that are candidates in both datasets are highlighted in purple. Candidates of both datasets are enriched within the region spanned by *3R Payne.* (C) Significant gene ontology (GO) terms based on differentially expressed genes among karyotypes at 18°C with Benjamini-Hochberg-adjusted *p*-values < 0.05. Also see supplementary table S5 (Supplementary Material online) and Materials and Methods for further details.

Similar to other *D. melanogaster* inversions (*In(2L)t*, *In(3R)Mo*, *In(3R)K*; see Lavington and Kern 2017; Said et al. 2018), *In(3R)Payne* karyotype affected DE across the entire genome (at 18°C; see fig. 8B; supplementary table S5, Supplementary Material online). These ‘non-local’ effects on DE suggest that the *3R Payne* inversion exerts major *trans-* acting regulatory effects (cf. Said et al. 2018), which is also consistent with the significant enrichment of DEG for gene ontology (GO) terms related to regulation of gene expression (fig. 8C). Despite these genome-wide transcriptional effects, DEG were enriched within the region spanned by the inversion (108 and 540 genes in-and outside *3R Payne*, respectively; Fisher’s exact test (FET), *p* < 0.001). By contrast, we failed to find enrichment for effects of temperature (459 and 5382 genes in-and outside *3R Payne*, respectively; FET, *p* = 0.75). Beyond DEG involved in regulating expression, GO analysis revealed that the inversion polymorphism also affects the expression of genes involved in growth, development, and reproduction (fig. 8C), as might be expected given the multifarious effects of *3R Payne* on fitness traits such as body size, survival upon starvation, cold shock survival and lifespan (Rako et al. 2006; Kapun et al. 2016b; Durmaz et al. 2018).

Since inversions can have a large impact on the expression of genes in the breakpoint regions (Lavington and Kern 2017; Said et al. 2018), we also asked whether the 108 DEG within the inverted region might be enriched in the breakpoint regions (breakpoints plus a region of up to ± 2 Mb proximal and distal to each breakpoint): there was no evidence for an uneven distribution of DEG as compared to expectations based on non-candidate genes (FET; *p* = 0.83). Given that *3R Payne* affects DE inside the inversion body as well as across the entire genome, variants in the breakpoints cannot fully account for the transcriptional effects of the inversion. These results agree well with the conjecture that inversions such as *In(3R)Payne* affect gene expression as a consequence of linked allelic variation maintained by selection for suppressed recombination (see Said et al. 2018).

To identify links between allelic variation and DEG with respect to karyotype, we compared genomic and transcriptomic candidates (supplementary table S3, supplementary table S7, Supplementary Material online). We first quantified the amount of overlap between karyotypic DE at 18°C (FIFS18 = Florida inverted vs. Florida standard reared at 18°C) and gene-wise *F*_ST_ without applying significance thresholds since arbitrary thresholds might constrain the ability to identify overlaps. Using rank-rank hypergeometric overlap tests (RRHO; Cahill et al. 2018) applied to all genes ranked either by *F*_ST_ or by DE, we found that only genes with high *F*_ST_ values exhibited highly significant overlap with strongly differentially expressed genes (fig. 9A). This analysis identified a core set of 86 overlapping genes (see top right corner of fig. 9A) which are all located within *In(3R)Payne* or in close proximity to it (fig. 8B; supplementary table S3, supplementary table S7; Supplementary Material online). Similar results were obtained when repeating the analysis with the data based on DE between karyotypes irrespective of rearing temperature (FIFS = Florida inverted vs. Florida standard; see supplementary fig. S9, Supplementary Material online). This provides further evidence that allelic variation inside the genomic region spanned by the inversion has a major functional impact on patterns of gene expression (cf. Said et al. 2018).

**Fig. 9.**
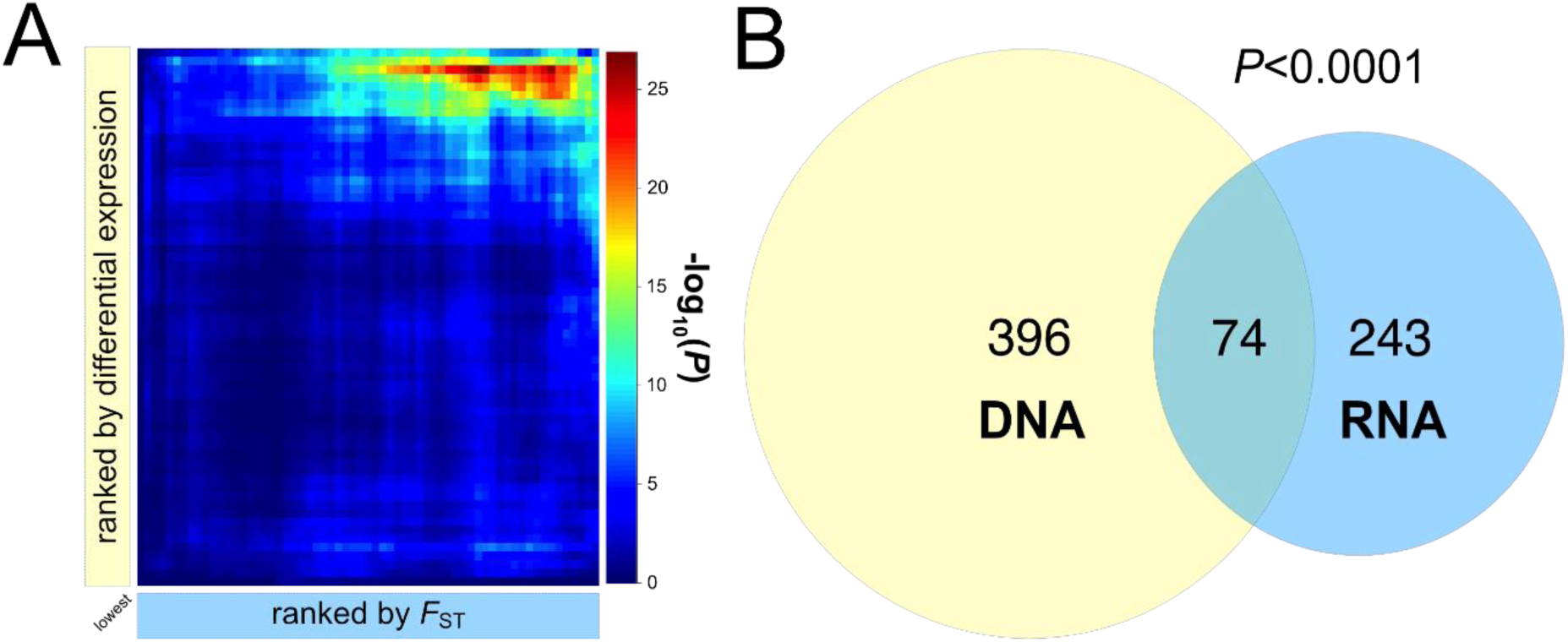
Overlap between genomic and transcriptomic candidate loci associated with *In(3R)Payne.* (A) Summary of results of rank-rank hypergeometric overlaps (RRHO). The dark red area indicates the highly significant overlap between genomic and transcriptomic candidates. A core set of 86 candidate loci located in the top right corner of the heatmap in (A) is tightly clustered inside the inversion or in close proximity to it (inversion highlighted in cyan). (B) Significant overlap between genomic candidates (based on candidate SNPs with *F*_ST_ ≥ 0.9 between karyotypes in Florida; in light yellow) and transcriptomic candidates (based on significant differential expression between inverted and standard karyotypes from Florida reared at 18°C = FIFS18; in light blue); *p*-value estimated using *SuperExactTest* in *R*. Also see supplementary figure S7, supplementary table S3 and supplementary table S7 (Supplementary Material online).

We subsequently focused on 470 candidate SNPs on *3R* with *F*_ST_ ≥ 0.9 between inverted and standard arrangements in Florida and 317 transcriptomic candidates with significant DE between karyotypes at 18°C (FIFS18). Comparison of these two groups of candidates revealed a significant overlap, comprising 74 genes (*SuperExactTest*; expected overlap: 37 genes; *p* < 0.001; fig. 9B; supplementary table S3, supplementary table S7, Supplementary Material online). Similarly, when considering 55 candidates exhibiting DE with respect to karyotype irrespective of developmental temperature (FIFS), we found a significant overlap of 19 genes (*SuperExactTest*; expected overlap: 4 genes; *p* < 0.001; supplementary fig. S9; supplementary table S3, supplementary table S7, Supplementary Material online). Although neither the 74 nor the 19 overlapping candidate loci were enriched for GO terms, several of them have well-known biological functions (supplementary table S3, Supplementary Material online; also see gene information on FlyBase at http://flybase.org/).

A comprehensive database of *In(3R)Payne*-associated candidate loci, based on our genomic, transcriptomic and overlap analyses, is provided in supplementary table S3 (Supplementary Material online). In addition to listing many novel candidates, this dataset contains and corroborates numerous genes previously associated with either latitudinal clinality and/or with *In(3R)Payne* (Hoffmann and Weeks 2007; Kolaczkowski et al. 2011; Fabian et al. 2012; Chen et al. 2012; Zhao et al. 2015; Kapun et al. 2016a, 2016b, 2020). These candidates include several loci with established mutant effects on fitness-related traits (e.g., size, reproduction, lifespan, stress resistance; cf. Kapun et al. 2016b; Durmaz et al. 2018; Kapun and Flatt 2019). Our database of candidates associated with *3R Payne* thus provides rich grounds for future work aiming to dissect the genetic architecture of this balanced inversion polymorphism.

## Conclusions

Here we have sought to refine our understanding of the adaptive nature of a common cosmopolitan chromosomal inversion polymorphism in *D. melanogaster*, *In(3R)Payne*, on four continents: in its ancestral African range and in derived populations in Europe, North America and Australia. Based on our population genomic and transcriptomic analyses, we offer the following conclusions and conjectures:

(1) Our data confirm that the *3R Payne* polymorphism is monophyletic, consistent with a single mutational origin in Africa at least ∼129 kya (see Corbett-Detig and Hartl 2012). Despite some genetic (geographical) divergence both within inverted and standard chromosomes among continents, inverted arrangements always cluster together, independently of their geographical provenance, and the same is true for standard arrangements.
(2) Phylogenetic analysis and patterns of divergence and LD support a scenario whereby differentiation between inverted and standard chromosomes worldwide is due to ancestral variants that differentiate the two karyotypes. This interpretation is supported by (1) significant sharing among continents of loci that are strongly differentiated between the karyotypes, both in the breakpoint regions and the center of the inversion, and (2) an absence of pronounced genetic divergence among inverted chromosomes from different continents. Analyses of inversion-specific alleles that are nearly or completely fixed in non-African populations within the African population sample suggest that the inversion has captured adaptive alleles in its ancestral range prior to the inversion having migrated out of Africa and become cosmopolitan.
(3) Patterns of nucleotide variability, genetic divergence and LD are consistent with (potentially long-term) balancing selection maintaining the inversion polymorphism (cf. Charlesworth 2023), but the exact type of balancing selection remains to be elucidated. Given its intermediate frequency in low-latitude populations and its absence in high-latitude locales around the world, *3R Payne* appears to have spread out of its ancestral tropical range and then become assorted by spatially varying selection in a parallel manner, causing the formation of similar clines on several continents (Kapun and Flatt 2019). This scenario is consistent with theoretical expectations suggesting that inversion frequencies can be maintained by balancing selection at local equilibria that change clinally (Faria et al. 2019); this could promote inversion polymorphism across large geographical scales and lead to parallel, stable large-scale clines (Westram et al. 2022).
(4) Our results and previous work (cf. Kapun and Flatt 2019) suggest that *3R Payne* is involved in parallel or ‘global’ (species-wide; cf. Booker et al. 2021) adaptation to similar latitudinal gradients around the world. It is noteworthy in this context that in Ethiopia *3R Payne* occurs at much lower frequency in a cold high-altitude locale as compared to a warm low-altitude habitat (Pool et al. 2017). Similarly, Aulard et al. (2002) found negative (albeit non-significant) correlations between *In(3R)Payne* frequency and altitude in African populations. Because *3R Payne* does typically not fix under warm conditions but appears to be selected against under cool conditions, it is an intriguing possibility that the loci captured by the inversion provide some form of balancing (e.g., negative frequency-dependent) selection independent of temperature yet happen to render it less tolerant to cool temperatures.
(5) RNA-seq analyses of inverted and standard chromosomes in a sample from North America (Florida) reveal pronounced effects of inversion karyotype on gene expression that depend on developmental temperature: expression levels are higher for inverted chromosomes at low temperature, perhaps due to a loss of buffering or compensatory plasticity (cf. Huang et al. 2022) and consistent with the notion that *3R Payne* is susceptible to cool conditions (see above; cf. Kapun et al. 2016a; Pool et al. 2017; Kapun and Flatt 2019).
(6) Although the inversion body is enriched for differentially expressed genes, the *3R Payne* inversion has pervasive genome-wide effects on gene expression, consistent with *trans-* acting regulatory effects. Functional effects of this inversion are thus unlikely explained by lesions at the breakpoints alone. Together with analyses of divergence and LD, our results support the idea that *3R Payne* maintains non-random associations among adaptive loci (Said et al. 2018). Yet, whether the linked loci are subject to epistatic balancing selection or to another selective mechanism is an open question. Likewise, the precise identity of the adaptive loci associated with the inversion remains unknown – our database of candidate loci might serve as a fruitful starting point for addressing this important question in future work.

## Materials and Methods

### Fly strains and their maintenance

To investigate karyotype-specific patterns of genetic variation and differentiation between karyotypes we established isofemale lines from populations in North America (Homestead, Florida, and Bowdoin, Maine, collected by Paul Schmidt) (see supplementary table S1, Supplementary Materials online). Lines were maintained under standard laboratory conditions (25°C; 60% relative humidity, 12h:12h light:dark cycle). We karyotyped these lines for six major chromosomal inversion polymorphisms (*In(2L)t, In(2R)NS, In(3L)P, In(3R)C, In(3R)Mo*, *In(3R)Payne*; see Lemeunier and Aulard 1992) using codominant PCR markers following the approach of Corbett-Detig et al. (2012). By combining PCR-based karyotyping and experimental crosses we generated lines that were isochromosomal for the third chromosome as described in Kapun et al. (2016b). In brief, we crossed wild-type males to females carrying a compound (second and third chromosome) balancer (*SMB6*; *TM6B*; Bloomington *Drosophila* Stock Center [BDSC] stock #5687) in an *ebony* (*e*^1^) mutant background. F1 offspring, which were heterozygous for the wild-type chromosome and the balancer, were visually selected based on the dominant *tubby* (*Tb*^1^) marker mutant phenotype and backcrossed to the balancer strain to amplify the wild-type chromosomes. Using PCR markers (Matzkin et al. 2005; Corbett-Detig et al. 2012) we determined the karyotype status of successfully isolated lines with respect to *In(3R)Payne* (see Kapun et al. 2016b for details). Whenever possible we selected against the balancer to establish isochromosomal lines. However, for a subset of standard and inverted chromosomes we failed to obtain homokaryons, likely due to recessive homozygous lethal alleles segregating among these chromosomes – this is not surprising given that typically ∼30-55% of wild third chromosomes are homozygous lethal (Simmons and Crow 1977; Mukai and Nagano 1983; our unpublished observations). In these cases, isolated wild-type chromosomes were propagated as heterozygotes over the balancer chromosome. During the propagation of the compound balancer line used for isolating wild third chromosomes, we occasionally observed that the visual marker of the second chromosome balancer (*SM6B*; *Duox*^Cy^) segregated independently of the visual marker of the third chromosome balancer (*TM6B*; *Tb*^1^). Since we had not consistently selected for both visual markers during the isolation process, we could not rule out that the wild-type second chromosomes might occasionally have recombined with those of the lab strain. We therefore excluded sequencing data from all chromosomal arms other than *3L* and *3R* from downstream genomic analyses.

### Sample preparation for DNA re-sequencing

We generated single-individual (phased) sequencing data using a subset of North American lines from Florida and Maine (see supplementary table S1, Supplementary Material online) to investigate patterns of linkage disequilibrium (LD) and haplotype structure with respect to karyotype and geographic origin. To obtain phased haploid sequencing data, we employed a ‘hemiclone’ approach, as described in Kapun et al. (2014) (supplementary fig. S10; Supplementary Material online). To this end, we crossed females from isochromosomal lines, or from strains with wild third chromosomes maintained over the balancer, to males of a highly inbred, inversion-free isofemale reference strain from Nigeria (line NG9 from the *Drosophila* Population Genomics Project [DPGP]; Lack et al. 2015). For each cross, we sequenced a single F1 hemiclonal male (supplementary fig. S10; Supplementary Material online). To bioinformatically discriminate between wild-type alleles and alleles segregating in the paternal NG9 reference (i.e., ‘bioinformatic phasing of alleles’) we pool-sequenced all reference strain males used for the crosses as a single pool (also see below).

### DNA extraction and library preparation

For each of the DNA libraries, we jointly homogenized whole tissue by bead beating (Zirconia beads; ⌀1.2 mm; 2 x 30 min at 6500 rpm) and incubated the homogenate in lysis buffer (100 mM Tris-Cl, 100 mM EDTA, 1% SDS, 1 mg/ml Proteinase K) for 30 min at 56°C and 30 min at 70°C. The lysate was treated with RNAse A (3 mg per 250 μl aliquot) at 37°C for 30 min prior to adding 39 μl of 8 M potassium acetate, followed by another incubation step for 30 min on ice to precipitate protein. After centrifugation at 14,000 rpm for 15 minutes, we mixed the supernatant with one volume of phenol-chloroform-isoamyl alcohol (ratio 25:24:1). The aqueous phase was further washed with 0.75 volume of chloroform prior to precipitation of DNA by adding 3 volumes of ice cold 100% ethanol. After incubation at 4°C for 2h, followed by centrifugation at 14,000 rpm for 15 minutes, we washed the pellet with 70% ethanol, dried it at room temperature and then resuspended the DNA in 50 μl of TE buffer. gDNA libraries of each sample were sheared using a Covaris instrument (Duty cycle 10%, intensity 5, cycles/burst 200, time 50s) and prepared for paired-end sequencing at the Lausanne Genomic Technologies Facility (GTF), using the Illumina TruSeq Nano Library preparation kit (Illumina, San Diego, USA). Samples were sequenced on a HiSeq 2000 Illumina Sequencer to 100 bp paired-end reads.

### Differential gene expression assays with RNA-seq

Given that *In(3R)Payne* is involved in climate adaptation (e.g., Kapun et al. 2016a, 2020, Kapun and Flatt 2019), and given that some the inversion’s phenotypic effects depend on growth temperature (Durmaz et al. 2018), we sought to examine the effects of *3R Payne* karyotype and/or of developmental temperature on gene expression and to identify potential candidate transcripts / genes associated with *In(3R)Payne.* To do so, we performed RNA-seq assays on isochromosomal lines carrying either the inverted or standard arrangement from Florida, where the inversion is polymorphic, and on lines carrying the standard arrangement from Maine, where the inversion is absent (see Kapun et al. 2016b for details of sampling locations; see Durmaz et al. 2018 for details of isochromosomal lines; also see supplementary table S1, Supplementary Material online). Each of these three groups was replicated 9-fold and exposed to two developmental temperatures (18°C, 25°C; see below) to account for interactions between genotype (karyotype) and environment (temperature). Prior to sampling for transcriptomic assays, flies were kept under common garden conditions (∼21°C, ∼50% relative humidity; 10h:14h L:D) for three generations. The experimental generation was reared at two growth temperatures during development until 5-7 days of adulthood (18°C vs. 25°C, 12:12h LD, 60% relative humidity, on a cornmeal/ sugar/ yeast/agar diet). Total RNA was extracted from 5-7 days-old snap-frozen adult females from each isochromosomal line, with each sample being prepared from 5 individuals (3 karyotypes x 2 temperatures x 9 isochromosomal lines = 54 samples) using the MagMAX™-96 Total RNA Isolation Kit (ThermoFisher Scientific, Waltham, MA, USA) on a MagMAX™ Express Magnetic Particle Processor (ThermoFisher Scientific, Waltham, MA, USA), following the manufacturer’s protocol. Prior to library preparation RNA quality was measured using Fragment Analyzer (Advanced Analytical) analysis. Single-end 101 bp long reads were sequenced on a Illumina HiSeq 2000 sequencer, following library preparation using the TrueSeq stranded library preparation kit. For details of bioinformatic analyses of these RNA-seq data see below.

### Bioinformatics pipeline for genomic analyses

The bioinformatics pipeline used for our population genomic analyses (see below for details), including scripts, is available here: https://github.com/capoony/In3RPayne_PopGenomics.

### Mapping pipeline

FASTQ reads from DNA and RNA sequencing data were examined for sequencing quality with FASTQC (v.0.10.1; http://www.bioinformatics.babraham.ac.uk/projects/fastqc/) and then trimmed and filtered with cutadapt (v.1.8.3; Martin 2011) to remove low-quality bases (base quality ≥ 18; sequence length ≥ 75 bp) and sequencing adapters. For DNA sequencing data, we only retained read pairs for which both reads fulfilled our quality criteria after trimming for mapping with bbmap (v.0.7.15; Li 2013) with default parameters against a compound reference genome consisting of the genomes of *D. melanogaster* (v.6.12) and common pro-and eukaryotic symbionts of *Drosophila*, including *Saccharomyces cerevisiae* (GCF_000146045.2), *Wolbachia pipientis* (NC_002978.6), *Pseudomonas entomophila* (NC_008027.1), *Commensalibacter intestine* (NZ_AGFR00000000.1), *Acetobacter pomorum* (NZ_AEUP00000000.1), *Gluconobacter morbifer* (NZ_AGQV00000000.1), *Providencia burhodogranariea* (NZ_AKKL00000000.1), *Providencia alcalifaciens* (NZ_AKKM01000049.1), *Providencia rettgeri* (NZ_AJSB00000000.1), *Enterococcus faecalis* (NC_004668.1), *Lactobacillus brevis* (NC_008497.1), and *Lactobacillus plantarum* (NC_004567.2), to avoid paralogous mapping. We filtered mapped reads for a mapping quality ≥ 20 and used Picard (v.2.17.6; http://picard.sourceforge.net) to remove duplicate reads and re-aligned sequences flanking insertions-deletions (indels) with the Genome Analysis Toolkit, GATK (v3.4-46; McKenna et al. 2010).

### Variant calling in DNA sequencing data

We combined mapped reads in BAM file format from each of the sequenced F1 hybrid individuals and from the sequenced pool of sires into a single mpileup file using samtools mpileup (v.1.3; Li *et al*. 2009) without base quality recalibration (parameter-B). Next, we reconstructed the identity of the maternal wild-type allelic state (‘bioinformatic phasing of alleles’) by contrasting polymorphisms present in the F1 larvae with the reference alleles from the sires based on the bioinformatics pipeline described in Kapun *et al*. (2014). We only considered positions that were homozygous in the reference pool (minimum minor allele frequency < 10%) and retained wild-type alleles with a minimum count of 10 across all sequenced F1 individuals. To avoid false positives, we excluded alleles whose counts fell outside the limits of a 90% binomial confidence interval based on an expected frequency of 50% at a heterozygous site in a given diploid F1 library. We further excluded positions with either (1) minimum coverage < 15 to reduce false negatives due to large sampling errors or (2) maximum coverage > the 95^th^ coverage percentile for the corresponding sample and chromosome to avoid false positives due to paralogous mapping. For positions with more than one wild-type allele, we only considered the most frequent allele.

### Sequencing data availability

All newly generated sequencing data are available under NCBI BioProject ID PRJNA928565 (http://www.ncbi.nlm.nih.gov/bioproject/928565).

### Additional sequencing data from other continents

To complement the above-mentioned sequencing data, we used previously published sequencing data from *D. melanogaster* lines from Africa, Europe and Australia, all with known inversion karyotype (see supplementary table S1, Supplementary Material online):

(i) African data. The African strains were collected in Siavonga (Zambia) and sequenced as haploid embryos to obtain fully phased sequences; they were bioinformatically karyotyped for various inversions as part of the DPGP (*Drosophila* Population Genomics Project) resource (see Lack et al. 2015, 2016). We focused on 21 lines that are known to segregate *In(3R)Payne* and randomly selected an equal number of strains with standard arrangement third chromosomes. Consensus sequence files were downloaded from the DGN (*Drosophila* Genome Nexus) website (http://www.johnpool.net/genomes.html), filtered for polymorphic sites and merged into a single VCF file using custom-made software. Genomic coordinates from *D. melanogaster* reference v.5 were converted to v.6.
(ii) European data. We used phased sequencing data from wild-type strains collected in Póvoa de Varzim in Portugal (Kapun et al. 2014; Franssen et al. 2015). In addition to 7 strains carrying *In(3R)Payne,* we randomly picked an equal number of strains with standard arrangement on the third chromosome and obtained the genomic data from Dryad (http://doi.org/10.5061/dryad.403b2). In addition to this European low-latitude sample from Portugal, we integrated phased sequencing data from 14 non-inverted strains from a high-latitude population in Umeå (Sweden) into our analyses, which were sequenced as haploid embryos (Kapopoulou et al. 2020), similar to the African samples mentioned above.
(iii) Australian data. Sequence data for the Australian continent were obtained from Rane et al. (2015) who had investigated population samples that approximate the endpoints of the latitudinal cline along the Australian east coast and sequenced these samples with reduced library representation RAD-tag sequencing. All 55 strains from Innisfail (tropical Queensland) (19 strains carrying *3R Payne* plus 18 carrying the standard arrangement) and Yering Station (temperate Victoria) (18 standard lines) had been screened by the authors for *3R Payne* using PCR markers (Rane et al. 2015). We obtained a VCF file containing high-confidence SNPs for all lines from Dryad (https://doi.org/10.5061/dryad.5q0m8) and converted genomic coordinates from *D. melanogaster* reference v.5 to v.6 prior to downstream analyses. Note that this dataset does not include genomic information for the first 12 million bp on chromosome arm *3R*.

### Re-analysis of DNA sequences of Australian isochromosomal lines

Rane et al. (2015) reported patterns genetic differentiation and LD with respect to *In(3R)Payne* in Australia that deviate from observations based on single-individual sequencing data of African flies (Corbett-Detig and Hartl 2012) and pool-seq data from North American flies (Kapun et al. 2016a). In contrast to the studies of Corbett-Detig and Hartl (2012) and Kapun et al. (2016a), Rane et al. (2015) found that Australian flies from tropical Queensland (where the polymorphism is segregating) do not exhibit elevated genetic differentiation between inverted and standard karyotypes within the genomic region spanned by *In(3R)Payne.* Yet, these authors found a pattern of strong, highly localized divergence between flies from Queensland and temperate flies from Victoria (where the inversion is very rare or absent), irrespective of *In(3R)Payne* karyotype. To explore why the patterns in the Australia data might differ from those observed in Africa and North America, we compared the 55 Australian libraries (Queensland: 19 inverted karyotypes, 18 standard karyotypes; Victoria: 18 standard karyotypes) to high-confidence sequencing data from 42 isogenic lines from Siavonga (Zambia, Africa; see above) which had previously been characterized for the presence or absence of *In(3R)Payne* (Lack et al. 2015, 2016). Our goal was to use these data to determine whether Australian and/or African lines cluster according to their *In(3R)Payne* karyotype and/or their geographic origin. Because *3R Payne* is of monophyletic African origin (Corbett-Detig and Hartl 2012, and analyses herein), we expected to find marked clustering of inverted chromosomes of African and Australian origin. Since gene flux due to double crossing overs is strongly suppressed or absent between the karyotypes in the breakpoint regions, we focused on 240 and 262 SNPs that were polymorphic both in Africa and Australia, respectively, and which were located within 200,000 bp around the proximal and distal breakpoints. We used custom-made software to combine and convert the allelic data from African and Australian lines to the NEXUS file format and calculated unrooted phylogenetic networks based on the Neighbor-Net inference method (see below). Since the karyotype-specific clustering of African strains and Australian lines from Queensland was inconsistent when using the karyotype classification of Rane et al. (2015) (supplementary fig. S4, Supplementary Material online), we used a panel of highly diagnostic, experimentally validated marker SNPs for *In(3R)Payne* (see Kapun et al. 2014 for details; also cf. Kapun et al. 2016a, 2020) to bioinformatically determine the karyotype status of the sequenced lines. Four of the 19 inversion-specific SNPs had sufficient coverage in the RAD-Seq data of most lines reported by Rane et al. (2015), thus allowing us to re-classify the karyotypes of the Australian lines. Notably, our new classification of karyotypes was highly consistent with the results of the clustering analysis of African samples. We therefore decided to use this new karyotype classification for all downstream analyses of the Australian data. Our analysis using inversion-specific marker SNPs also indicated that several Australian lines were not fixed for either the inverted or standard karyotype but appeared to be heterokaryotypic (supplementary table S8, Supplementary Material online). We thus excluded all apparently heterokaryotypic and/or ambiguous strains and only retained unambiguous homokaryotypes for downstream analyses.

### Phylogenetic relationships among *In(3R)Payne* karyotypes

Phylogenetic relationships among a total of 450 *D. melanogaster* strains from Africa, Europe, North America and Australia were analyzed with respect to *In(3R)Payne* based on a compilation of sequencing data from the above-mentioned sources (see supplementary table S1, Supplementary Material online). First, to investigate phylogenetic relationships among strains within the region spanned by *In(3R)Payne,* we analyzed 3766 SNPs located within the breakpoints of *In(3R)Payne.* Secondly, we reconstructed genome-wide patterns of phylogenetic relationships among these samples independent of chromosomal inversions by focusing on 4849 SNPs that were randomly drawn from the left and right arms of the third chromosome in 200 kb distance from the breakpoints of *In(3L)P* and *In(3R)Payne.* Using custom-made software, we combined and converted allelic data from all lines to the NEXUS file format and calculated unrooted phylogenetic networks based on the Neighbor-Net inference method (Bryant and Moulton 2004) with Splitstree (v.4.14.6; Huson 1998), using the Jukes-Cantor model for computing genetic distances. Importantly, unlike the Neighbor-Joining (NJ) method, Neighbor-Net can represent conflicting signals in the data, e.g., due to recombination (Bryant and Moulton 2004).

### Population genetic analyses

#### Analysis of nucleotide diversity and Tajima’s D

We quantified genetic variation using the software packages *vcftools* (v.0.1.16) to obtain SNP-wise estimates of nucleotide diversity *π* and Tajima’s *D* in samples with phased sequencing data. Because *vcftools* provides window-wise estimates based on total window size but does not account for positions in a given window that do not fulfill the same quality criteria as the polymorphic sites, we obtained average values in 100 kb non-overlapping windows using custom-made software. We first generated mask files where positions for which more than 50% of individuals did not fulfil heuristic quality criteria (based on minimum and maximum coverage, as defined above) were flagged with a ‘0’, whereas all other positions that passed were flagged with a ‘1’. We then calculated window-wise averages of *π* and Tajima’s *D* separately for inverted vs. standard chromosomes using the information in the mask files for population samples with phased sequencing data. To test for differences in genetic variation with respect to (1) geography, (2) karyotype and (3) genomic region we analyzed samples from Siavonga (Zambia, Africa), Póvoa de Varzim (Portugal, Europe), Homestead (Florida, USA) and Innisfail (Queensland, Australia). We considered all window-wise averages of *π* and Tajima’s *D* between positions *3R:*16,432,209 and *3R:*24,744,010 as being located ‘inside’ *In(3R)Payne.* To define a representative ‘control’ region ‘outside’ *of In(3R)Payne* we choose a random sample of equal size, composed of average estimates of *π* from *3L* and *3R* located outside the interval ranging from *3R:*14,232,209 to *3R:*26,744,010. To account for potential long-range effects of *In(3R)Payne,* we extended the actual length of the inversion by 2 Mb on both ends. Using *R* we performed a three-way analysis of variance (ANOVA) of the form *y*_i_ = *O* + *K* + *G* + *O* ´ *K* + *O* ´ *G* + *G* ´ *K* + *O* ´ *G* ´ *K*+ *ε*_i_, where *y*_i_ is the continuous dependent variable *π* or Tajima’s *D* in the *i*^th^ sample, *O* denotes the categorical factor ‘origin’ with four levels (Africa, Europe, North America, Australia), *K* represents the factor ‘karyotype’ with two levels (inverted, non-inverted) and *G* stands for the factor ‘Genomic region’ with two levels (in-vs. outside of the inversion), followed by all possible interactions, and where *ε* represents the error term. Based on the coefficients estimated from this model, we calculated planned contrasts using the *R* package *emmeans* to test for significant differences in genetic variation between karyotypes inside the inverted region. To search for a potential signature of balancing selection (as indicated by positive *D* values) we also estimated Tajima’s *D* for pooled samples of inverted and standard chromosomes (see supplementary fig. S2, Supplementary Material online). We note that estimating *D* for pools consisting of equal numbers of inverted vs. standard chromosomes (50:50 ratio; see supplementary fig. S2, Supplementary Material online) vs. *estimating D* using pools with the numbers of inverted vs. standard chromosomes being proportional to their population frequencies (not shown) did not make a qualitative difference.

#### Analysis of linkage disequilibrium (LD)

We estimated LD within and among karyotypes from low-latitude populations in Siavonga (Zambia, Africa), Póvoa de Varzim (Portugal, Europe), Homestead (Florida, USA) and Innisfail (Queensland, Australia) for which phased sequencing data were available. Squared allele frequency correlations (*r*^2^) (Hill and Robertson 1968) were calculated among pairs of 5000 randomly drawn SNPs on *3R* and between all polymorphic SNPs and *In(3R)Payne* using custom-made software as described in Kapun et al. (2014). Since the *r*^2^ statistic can be affected by large variance due to rare alleles, and because this might confound analyses of LD patterns (Hedrick 1987), we restricted analyses to SNPs with minor allele frequencies ≥ 0.1. To compare the decay of LD with physical distance we focused on 5000 SNPs located within the region spanned by *In(3R)Payne* and restricted analyses to pairwise *r*^2^ among SNPs within 100 kb distance. Following the approach in Remington et al. (2001) and Marroni et al. (2011), we used our LD estimates to fit the following equation from Hill and Weir (1988):

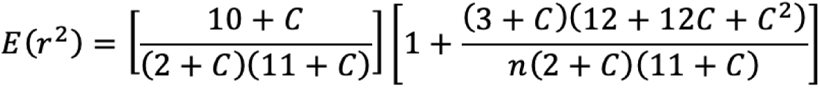

which allows modeling the expected reduction of *r*^2^ with physical distance. Here, *E*(*r*^2^) is the expected value of *r*^2^, *n* represents the sample size, and *C* is the product of the population-scaled recombination rate (*ρ* = 4*N_e_r*) and the distance in base pairs, which we estimated using nonlinear regression in *R.* For each population sample we employed the *R* function *nls* which fits a model based on nonlinear least squares across both karyotypes. Subsequently, we fitted the same model but additionally accounted for karyotype as a grouping factor using the function *nlsList* in the *R* package *nlme* (Pinheiro *et al*. 2021). To infer significant variation in the decay of LD as a function of *In(3R)Payne* karyotype we tested for differences in the goodness-of-fit of the two nested models by analysis of deviance, using the function *anova_nlslist* in the *R* package *nlshelper* (Duursma 2017).

#### Analysis of genetic differentiation

To quantify the amount of genetic differentiation between samples for single SNPs and for averages in 100 kb non-overlapping windows, we estimated pairwise *F*_ST_ for every SNP based on the method of Weir and Cockerham (1984), using *vcftools* (v.0.1.16). We first investigated how *In(3R)Payne* affects genetic differentiation within and among populations. We focused on population samples from the endpoints of latitudinal gradients in Europe, North America and Australia for which the low-latitude populations harbored *In(3R)Payne* at appreciable frequencies and for which the inversion was absent in high-latitude populations. Our LD analyses mentioned above revealed elevated LD inside inverted karyotypes, indicating that adjacent SNPs do not evolve independently. We thus compared average *F*_ST_ in 100 kb non-overlapping windows within the inversion breakpoints (‘inside’) to a similarly sized ‘outside’ set of average *F*_ST_ values that were randomly chosen from the third chromosome in 2 Mb distance from the breakpoints of *In(3L)P* and *In(3R)Payne,* as defined above. For the regions defined as ‘inside’ and ‘outside’ the inversion, and for each continent separately, we tested for differences in average pairwise *F*_ST_ values using the following comparisons as input data: (1) samples from the same low-latitude population with different karyotypes (factor level: ‘Karyotype’; e.g., Florida inverted vs. Florida standard [FIFS]), (2) samples with standard arrangement from different populations at the endpoints of a given continental latitudinal gradient (factor level: ‘Geography’; e.g., Florida standard vs. Maine standard [FSMS) and (3) samples with different karyotypes from different geographical populations within the same continent (factor l. evel: ‘Geography + Karyotype’; e.g., Florida inverted vs. Maine standard [FIMS]; i.e., pairwise *F*_ST_ estimates for which the effects of karyotype and geography might be confounded). To analyze these data we used one-way ANOVA of the form *y_i_* = *C* + *ε_i_*, where *y_i_* represents pairwise *F*_ST_ in the *i*^th^ genomic window, *C* is the categorical factor ‘pairwise comparison’ with three levels (‘Karyotype’, ‘Geography’, ‘Karyotype + Geography’) and *ε* represents the error term. To determine which of the three levels of *C* differ from each other we performed Tukey’s HSD post-hoc tests. The above-mentioned between-karyotype *F*_ST_ estimates (i.e., estimates of *F*_AT_, Charlesworth et al. 1997) were obtained for pools consisting of equal numbers of inverted vs. standard chromosomes (50:50 ratio); we note that estimating karyotypic divergence using pools with the numbers of inverted vs. standard chromosomes being proportional to their population frequencies did not make a qualitative difference (not shown). In addition to *F*_ST_, we also estimated genetic differentiation between karyotypes using *K*_ST_ (Hudson et al. 1992; also cf. Nei 1973 and Charlesworth 1998) – these analyses yielded qualitatively identical patterns as those for *F*_ST_ (results not shown). Finally, we also examined whether *F*_ST_ was elevated within the breakpoints of *3R Payne* when comparing inverted chromosomes between continents. We focused on SNPs shared across populations and calculated *F*_ST_ in 100 kb non-overlapping windows in all pairwise combinations across inverted chromosomes from Zambia, Portugal, Florida (USA) and Queensland (Australia) using *vcftools* (v.0.1.16). *F*_ST_ values were plotted against their genomic positions for all comparisons in *R* using the ggplot2 package.

#### Identification of candidate genes and SNPs associated with In(3R)Payne

To identify candidate genes and SNPs in the region spanned by by *In(3R)Payne,* we focused again on the samples from Siavonga (Zambia, Africa), Póvoa de Varzim (Portugal, Europe), Homestead (Florida, USA) and Innisfail (Queensland, Australia) and isolated SNPs positions that exhibited *F*_ST_ ≥ 0.9 between *In(3R)Payne* inverted and standard individuals within a given population; we considered all genes as candidate loci if at least one candidate SNP with *F*_ST_ ≥ 0.9 was located inside or within 2 kb proximity of the 3’ and 5’ ends of a given gene, since these regions harbour regulatory elements (Down *et al*. 2007; Nègre *et al* 2011). Long genes have a higher probability to harbor candidate SNPs by chance, and this might result in a bias towards gene ontology (GO) classes that are enriched for long genes. To account for this potential bias, we used *Gowinda* (Kofler and Schlötterer 2012) in order to test for overrepresentation of GO terms associated with karyotype-specific SNP candidates. *Gowinda* first generates an empirical null distribution of gene abundance in a given GO category based on a set of randomly chosen SNPs of equal size as the candidate set; *Gowinda* then estimates the significance of overrepresentation for each GO category and accounts for multiple testing by using Benjamini-Hochberg (BH) correction of *p*-values (Benjamini and Hochberg 1995). Next, we examined the extent to which candidate SNPs and genes are shared among continents. Using the *R* package *SuperExactTest* (Wang et al. 2015), which allows assessing the significance of intersections among multiple sets of similar data, we tested for overlaps between the sets of candidate SNPs and genes. Since *SuperExactTest* estimates predicted intersections based on the size of statistical background populations from which all sets are sampled, we only included SNPs that were polymorphic in all four datasets. Because the Australian samples were sequenced with RAD-Seq, only a limited number of SNPs could be recovered when comparing across all four populations (fig. 7). We thus excluded the Australian samples and performed genome-wide comparisons among the three remaining populations / continents. Based on the annotations assigned to each SNP with SNPeff (Cingolani *et al*. 2012), we tested for enrichment of candidates with Fisher’s exact test (FET) using a custom-made python script. We focused on the 8 most common SNP categories (i.e., intergenic_region, upstream_gene_variant, 5_prime_UTR_variant, intron_variant, synonymous_variant, missense_variant, 3_prime_UTR_variant and downstream_gene_variant). To build contingency tables for category-specific FETs, SNPs were classified as candidate vs. non-candidate and as belonging to a given category or not. To account for multiple testing *p*-values were Bonferroni-corrected.

### RNA-seq data analysis

Prior to mapping, we trimmed and filtered raw reads for a base quality ≥ 18 and read lengths ≥ 75 bp using *cutadapt*, as explained above. Next, we used *kallisto* (v.0.44.0; Bray *et al*. 2016) for pseudo-alignments of each library against the *D. melanogaster* transcriptome (v. 6.17, obtained from http://flybase.org/), using the following parameters: –l 101 (average fragment length = 101 bp); –s 10 (average standard deviation of fragment length = 10); –b 100 (number of bootstrapped samples = 100); ––rf-strand (reads are strand-specific, with the first read being reversed); ––single (reads are single-ended). We focused on gene-specific expression patterns and summed up all transcript-specific read counts for each gene using custom-made software following the approach in Soneson et al. (2015). We first transformed the raw absolute read counts to relative counts per million (cpm) and normalized data using the ‘trimmed means of the M-value’ (TMM) approach implemented in the *R* package *edgeR* (3.20.9) (Robinson et al. 2010, 2011). Lowly expressed or non-expressed genes were excluded from downstream analyses by removing genes with less than or equal to 2 CPM in more than or equal to 9 samples from the dataset. To identify differentially expressed genes affected by karyotype, developmental temperature or both, we fitted linear models to the expression data using the *R* package *limma* (Smyth 2005; Ritchie et al. 2015). By employing the *model*.*matrix* function of the *limma* package we set up a design matrix of the form: ∼*0* + *G*, where *0* indicates that the model is fitted without intercept and where *G* is a grouping factor with six levels (FI-18, FI-25, FS-18, FS-25, MS-18, MS-25) based on the geographic origin and karyotype of the samples (i.e., Florida inverted [FI], Florida standard [FS], Maine standard [MS]) and the two rearing temperatures (18°C, 25°C). To account for potential line effects, we included replicate line identity as a random effect nested in karyotype. Using the *voom* function of *limma*, precision weights were calculated for log_2_-CPM-transformed read counts to account for the relationship between mean and variance in RNA-Seq data when fitting linear models to expression data (Law et al. 2014). We used the *eBayes* function to improve the accuracy of gene-wise variance estimates by empirical Bayes moderation which integrates information on variation across all genes in the dataset. Based on the parameter estimates for each of the six levels of the grouping factor *G*, we calculated contrasts to test for the effects of karyotype, geography, and karyotype + geography, averaging across both temperatures. In addition, we calculated contrasts for the two developmental temperatures separately. We also employed contrasts to examine interactions between temperature and karyotype, geography and karyotype + geography. To account for multiple testing, we used BH *p*-value correction and only considered genes with a *p* < 0.05 to be differentially expressed. For each of the candidate gene lists obtained from differential expression analysis with *limma*, we tested for enrichment of specific GO categories using the *R* package *topgo* (Alexa and Rahnenführer 2009). After correcting significant sets of GO terms for hierarchical clustering using *GO-Module* (Yang et al. 2011), we visualized the remaining set of GO terms with REVIGO (Supek et al. 2011) and Cytoscape (Shannon et al. 2003). Enrichment of candidates according to their position relative to *In(3R)Payne* was tested by creating contingency tables based on candidate and non-candidate genes located either in-or outside the region spanned by the breakpoints of *In(3R)Payne* and using Fisher’s exact tests in *R*. We further compared our candidates to the candidates reported by Zhao et al. (2015), who had identified genes that exhibit differential expression between a low-latitude population in Panama and a high-latitude population in Maine at two rearing temperatures (21°C, 29°C), and tested for significant overlaps using *SuperExactTests* (Wang et al. 2015).

### Overlap between genomic and transcriptomic candidates

To refine our set of candidate loci associated with *3R Payne* we compared genomic and transcriptomic candidates. Genomic plus transcriptomic data were only available from North American populations (i.e., inverted and standard karyotypes from Florida; standard karyotypes from Maine). We tested for significant overlaps between *F*_ST_-based and differentially expressed candidate genes using *SuperExactTests* (Wang et al. 2015); as the background set for these analyses we only considered third chromosome genes identified in both datasets. Since the significance of overlaps across sets can be confounded by the choice of significance thresholds in the individual datasets, we also employed a comparison based on rank-rank hypergeometric overlaps of ranked gene lists using the *R* package RRHO (Plaisier et al. 2010). RNA-seq candidates were ranked based on adjusted *p*-values, whereas genomic candidates were ranked based on average *F*_ST_ of the 10% top most highly differentiated SNPs located within 2 kb proximity of a given candidate gene. RRHO tests for significant overlaps between gene lists are based on hypergeometric tests calculated while sliding across all possible thresholds in the two ranked lists. Besides visual representation of changes of significance with decreasing rank, RRHO allows to define an ‘optimally’ overlapping gene set. We used this ‘optimal’ set to test for enrichment of GO categories using *topGO* in the *R* package *limma*.

## Supporting information

STable S3

STable S4

STable S5

STable S6

STable S7

STable S8

STable S1

STable S2

## Acknowledgements

We are indebted to Brian Charlesworth for helpful comments on previous versions of the manuscript and for sharing unpublished results; to Mark Kirkpatrick for suggesting the analysis in figure 6; and to two anonymous reviewers for their valuable feedback. We also acknowledge the Lausanne Genomic Technologies Facility (GTF) and the Vital-IT Bioinformatics Facility at the University of Lausanne for sequencing and bioinformatics support. Our research was funded by the Swiss National Science Foundation (SNSF grants 31003A-182262, PP00P3_165836, PP00P3_133641/1 to TF), the Austrian Science Fund (FWF grant P32275 to MK), the Department of Ecology and Evolution at the University of Lausanne, and the Department of Biology at the University of Fribourg. TF also received support from a Mercator Fellowship of the German Research Foundation (DFG), held as an EvoPAD Visiting Professor at the Institute for Evolution and Biodiversity, University of Münster (Germany).

## Author contributions

Contributions following CRediT (Contributor Roles Taxonomy; https://casrai.org/credit): <u>MK</u>: Conceptualization, Data curation, Formal Analysis, Funding acquisition, Investigation, Methodology, Resources, Software, Validation, Visualization, Writing – original draft, Writing – review & editing; <u>ED</u>: Formal Analysis, Investigation, Methodology, Software, Validation, Visualization, Writing – original draft, Writing – review & editing; <u>TJK</u>: Funding acquisition, Writing – review & editing; <u>PS</u>: Resources, Writing – review & editing; <u>TF</u>: Conceptualization, Funding acquisition, Investigation, Project administration, Resources, Supervision, Validation, Writing – original draft, Writing – review & editing.

