## Supplementary material for "An Ancestral Balanced Inversion Polymorphism Confers Global Adaptation": Supp Figures

Running title: Genomics of a Balanced Inversion Polymorphism

<u>**Supplementary Material Online**</u>

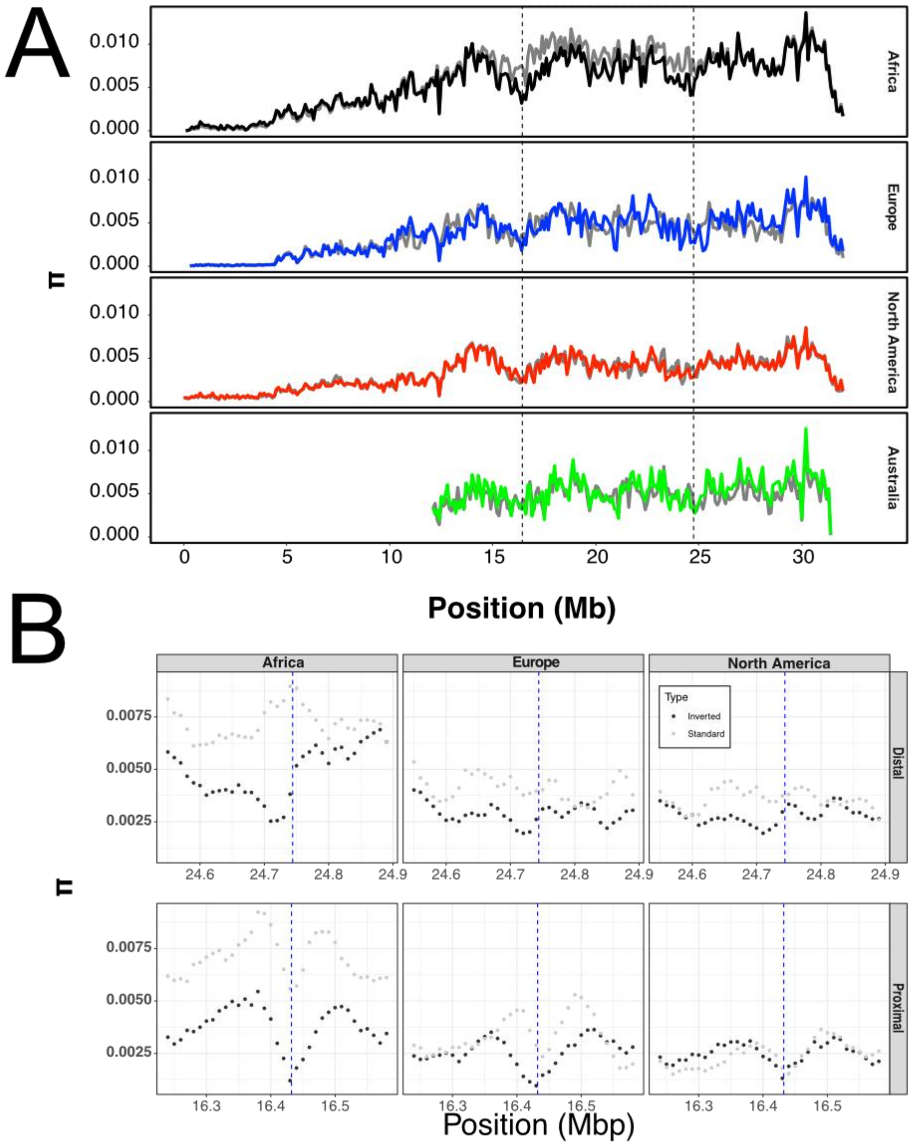


**Supplementary figure S1.** (A) Window-wise estimates of nucleotide diversity (*π*) along chromosomal arm *3R* in non-overlapping windows of 100 kb length. Estimates of standard arrangement chromosomes are shown in grey and estimates of inverted chromosomes in black, blue, red and green for samples from Africa, Europe, North America and Australia, respectively. The genomic position of *In(3R)Payne* is highlighted by a dashed black outline. Also see main text fig. 2A. (B) Zoom-in of the distal (top panel) and proximal (bottom panel) breakpoint regions, showing estimates of nucleotide diversity (*π*) averaged in 50 kb windows with 10 kb steps as grey dots for standard and as black dots for inverted chromosomes.

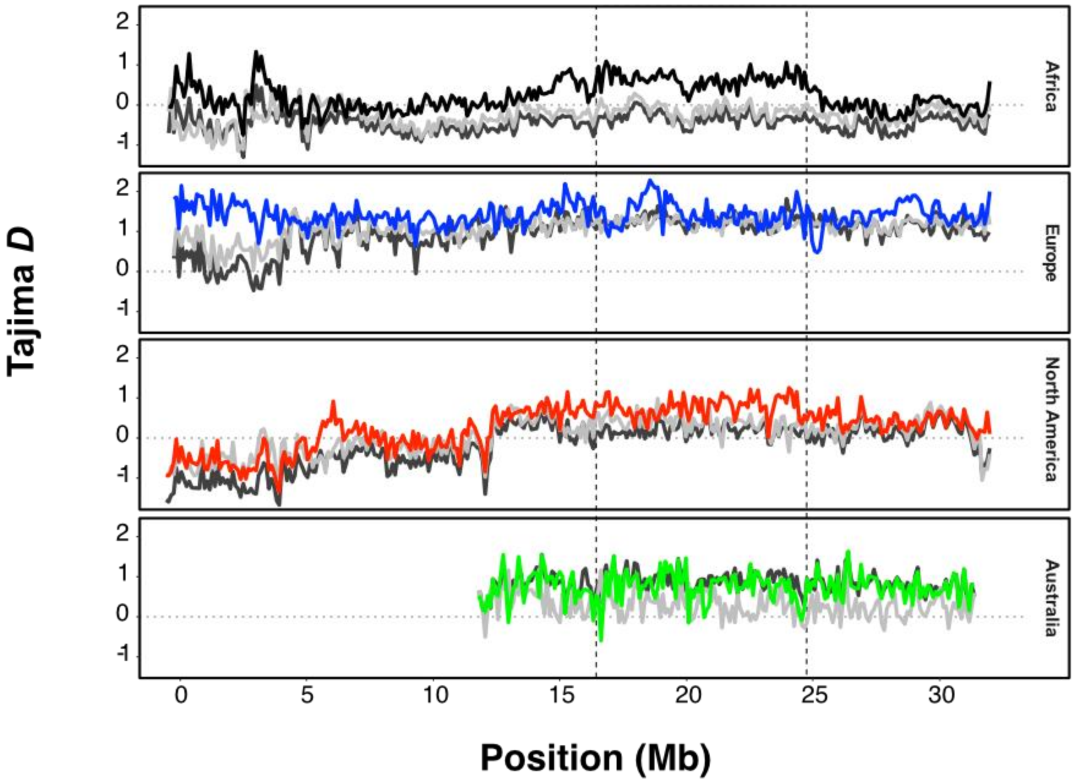


**Supplementary figure S2.** Window-wise estimates of Tajima's *D* along chromosomal arm *3R* in non-overlapping windows of 100 kb length, based on equal numbers of chromosomes for each karyotype and population. Estimates for all haplotypes pooled irrespective of karyotype are shown in dark grey;  for standard arrangements in light grey; and for inverted chromosomes in black, blue, red and green for samples from Africa, Europe, North America and Australia, respectively. The genomic region spanned by *In(3R)Payne* is delineated by vertical dashed lines; the vertical dotted line indicates Tajima's *D* = 0, i.e., value of *D* expected under standard neutrality. Also see main text fig. 2B.

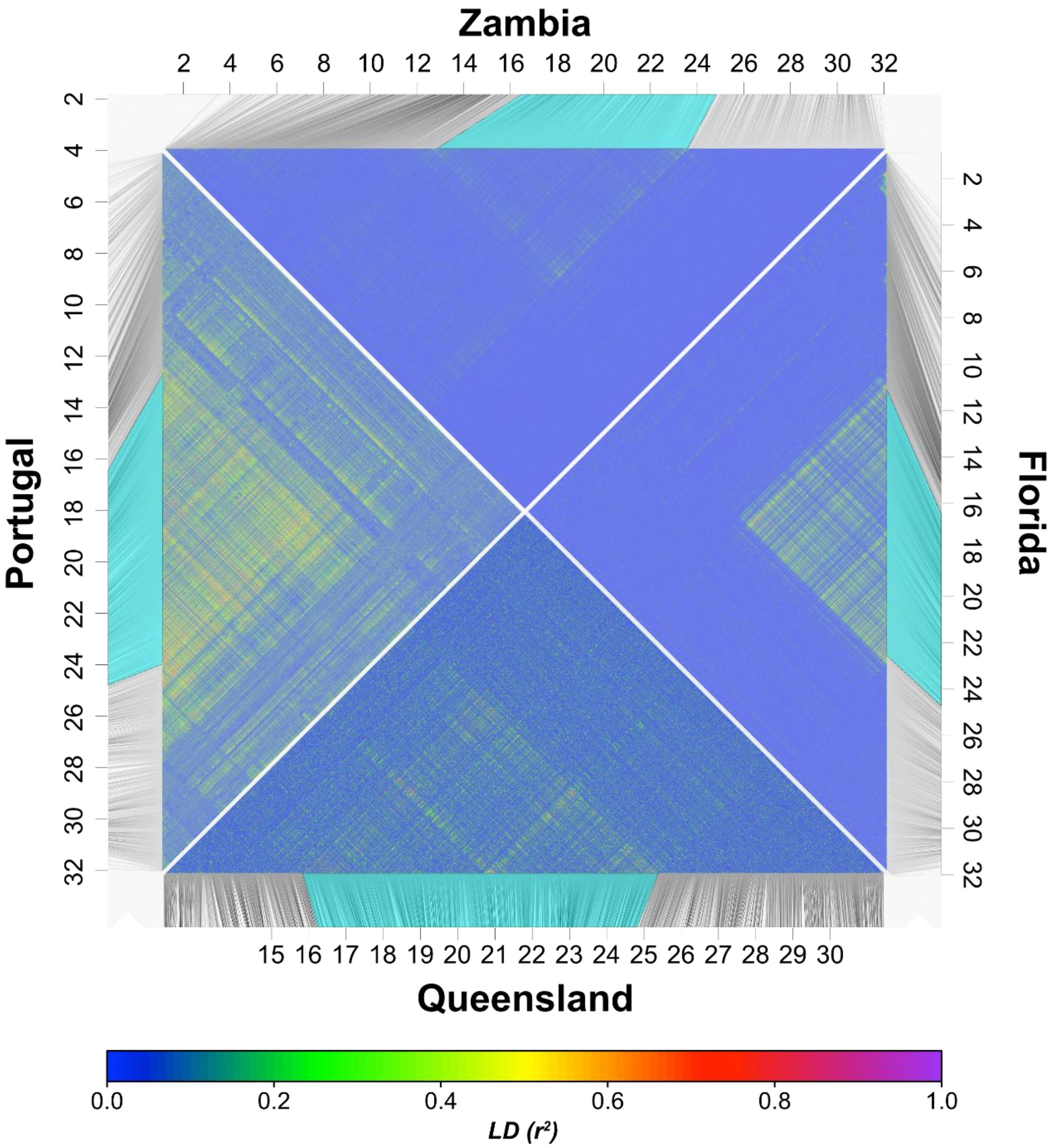


**Supplementary figure S3.** Pairwise values of $r^2$ as an estimate of linkage disequilibrium based on 5000 randomly drawn SNPs along chromosomal arm *3R* for samples from Africa (Zambia), Europe (Portugal), North America (Florida) and Australia (Queensland), respectively. All samples consisted of equal numbers of inverted and standard chromosomes. The areas highlighted in cyan represent the genomic region spanned by *In(3R)Payne.* Also see main text fig. 3.

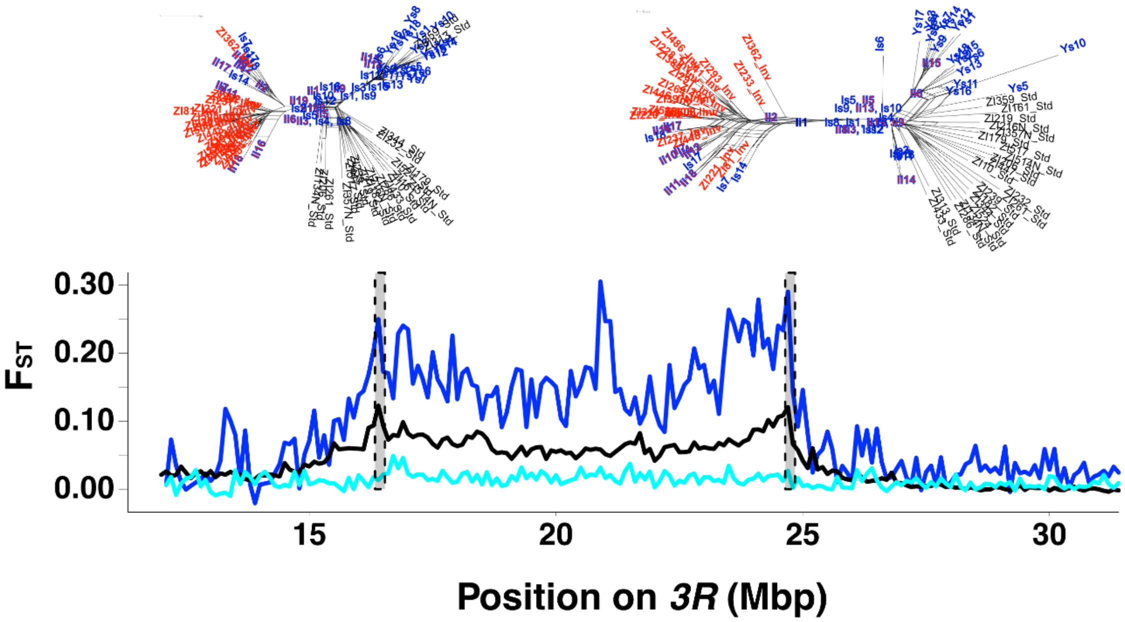


**Supplementary figure S4.** Reclassification of *In(3R)Payne* karyotypes for Australian samples from Rane et al. (2015). The two phylogenetic networks at the top are based on SNPs within 200kb genomic regions spanning the proximal breakpoint at *3R*:16,432,209 and the distal breakpoint at *3R*:24,744,010 of *In(3R)Payne* (indicated by dashed outlines in the bottom line plots), respectively. We focused on these regions because they should not be affected by double recombination; for phylogenetic analysis, we included all Australian samples from Queensland (Innisfail) and Victoria (Yering Station) from Rane et al. (2015), as well as 21 standard and 21 inverted samples from Siavonga (Zambia, Africa) for reference. Both networks indicate that several of the Australian samples from Victoria (symbols with blue fill and outline and marked with "Y"), previously classified as inverted arrangement (symbols with red fill and blue outline) by Rane et al. (2015), cluster clearly within the standard arrangement samples from Zambia (indicated in black). Conversely, several Australian samples from Queensland (symbols with blue fill and outline and marked with "I"), previously classified as standard arrangement (Rane et al. 2015), cluster within inverted Zambian samples (indicated in red). Together, our analyses strongly suggest that several of these samples were previously misclassified. The line plots at the bottom show values of $F_{ST}$ in 100 kb non-overlapping windows among karyotypes (1) according to the original classification of Rane et al. (2015) in cyan, (2) following our new classification based on the above-mentioned phylogenetic approach in blue, and (3) among African samples of different karyotype from Zambia for reference in black. Consistent with previous misclassification, $F_{ST}$ patterns based on our new classification reveal high divergence at the breakpoints and a massive central peak in the inversion body, a pattern that is highly consistent with similar data from other continents (see fig. 4 and supplementary fig. S5).

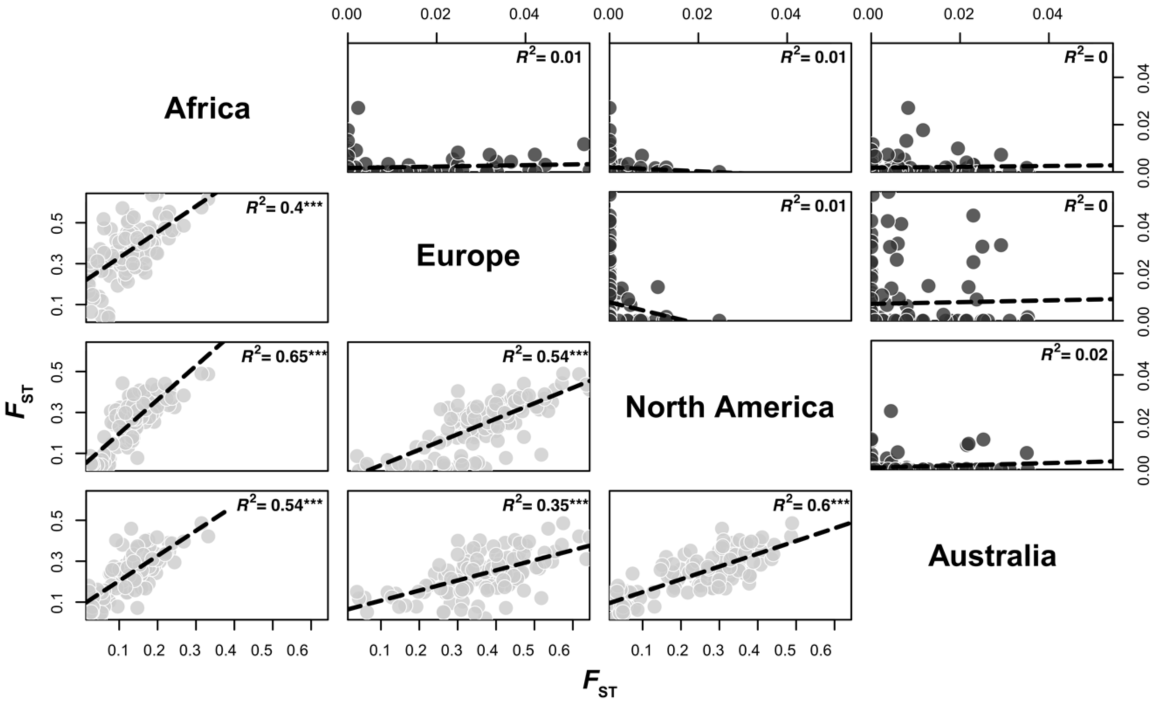


**Supplementary figure S5.** Pairwise correlations of $F_{ST}$ in 100 kb windows for samples from different continents. The subplots in light grey below the diagonal show correlations of $F_{ST}$ windows inside *3R Payne*; subplots in dark grey above the diagonal show similar analyses based on windows in the genomic region between *3L*:4,432,209 and *3L*:16,744,010, for comparison. $R^2$ values (coefficient of determination, proportion of variance explained) are shown in the top right corner of each plot; *** $p$ < 0.001 based on linear regressions.

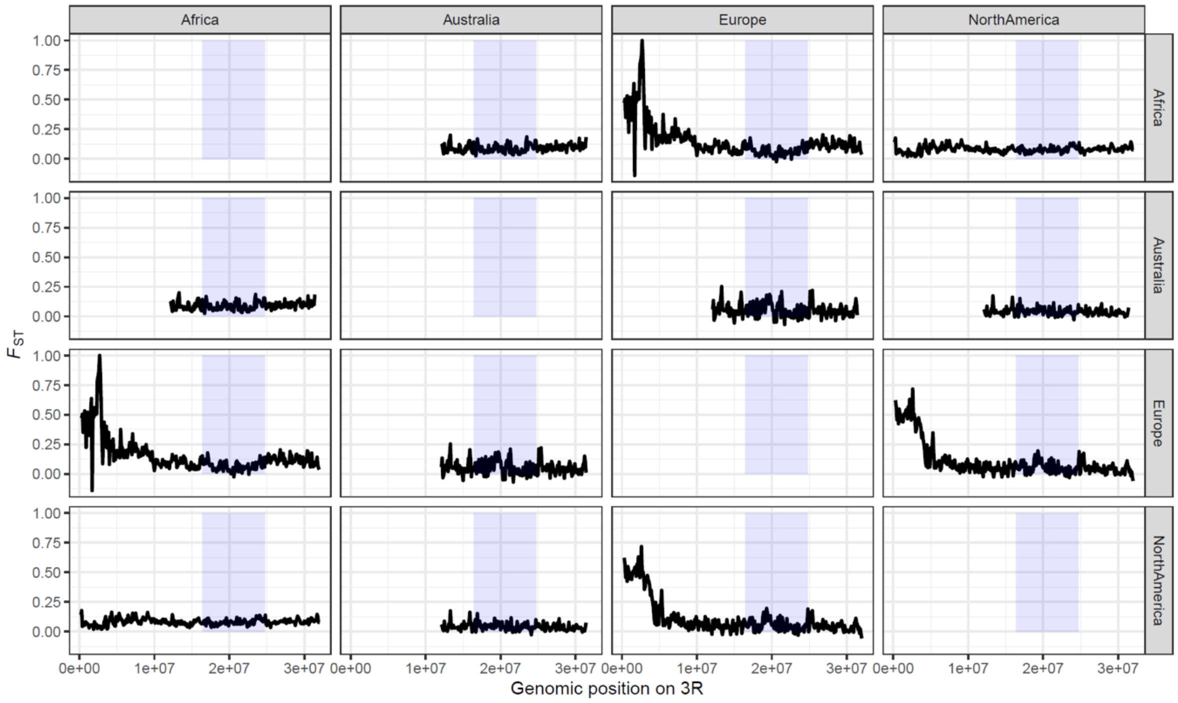


**Supplementary figure S6.** Pairwise $F_{ST}$ in 100 kb non-overlapping among inverted chromosomes carrying *In(3R)Payne* from different continents. The subplots below and above the diagonal are identical; the light blue boxes highlight the genomic region spanned by *In(3R)Payne*.

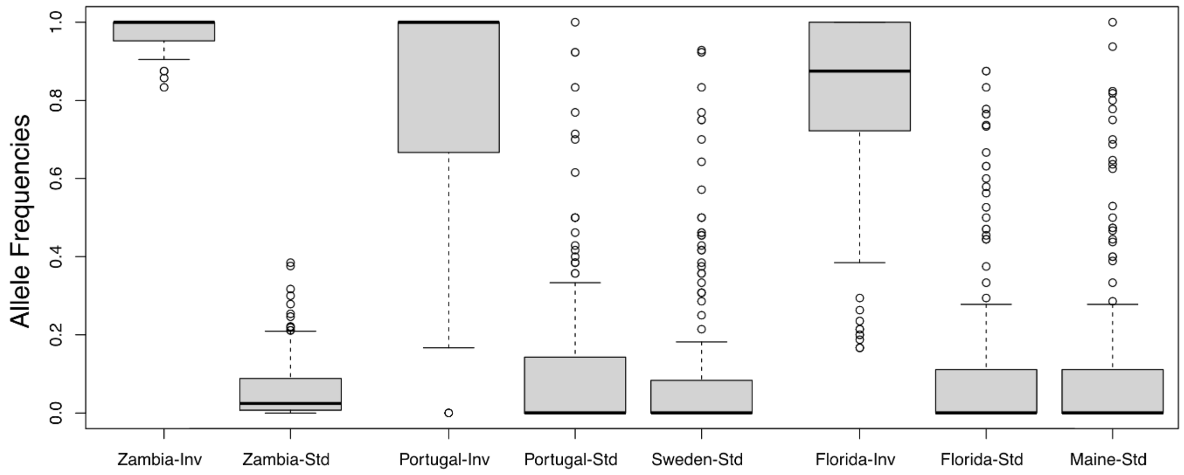


**Supplementary figure S7.** Allele frequencies of 'inversion-specific' SNPs in derived (non-African) populations that are characterized by $F_{ST}$ ≥ 0.9 between inverted and standard chromosomes based on the African population sample from Zambia (also see fig. 6A). In the North American (Florida) sample, for example, inverted chromosomes are also polymorphic for standard African alleles. This suggests that inversion-specific alleles in Florida were not acquired locally but that they are ancestral and predate the out-of-Africa expansion.

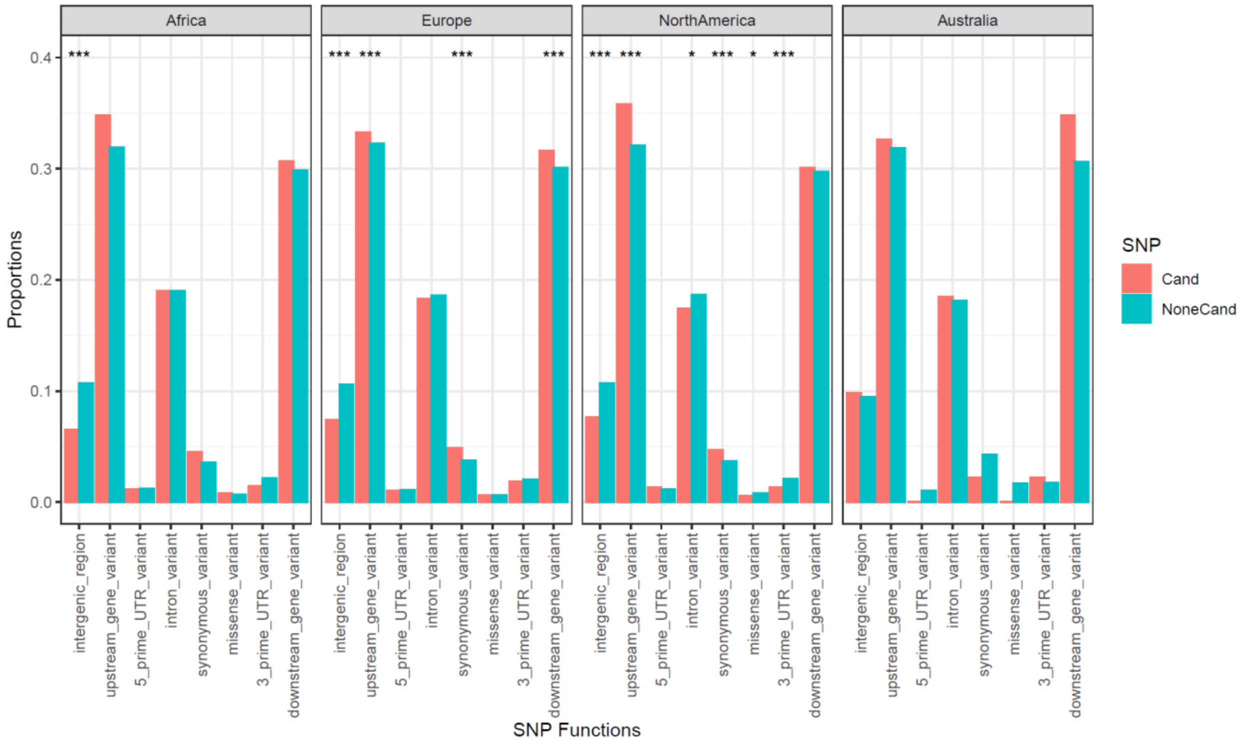


**Supplementary figure S8.** Bar plots showing the proportions of candidate (salmon) and non-candidate (turquoise) SNPs belonging to one of 8 major functional categories of genic positions. Significant proportional differences between candidates and non-candidates based on Fisher's exact tests are highlighted by asterisks above the corresponding bars. * $p < 0.05$; ** $p < 0.01$; *** $p < 0.001$.

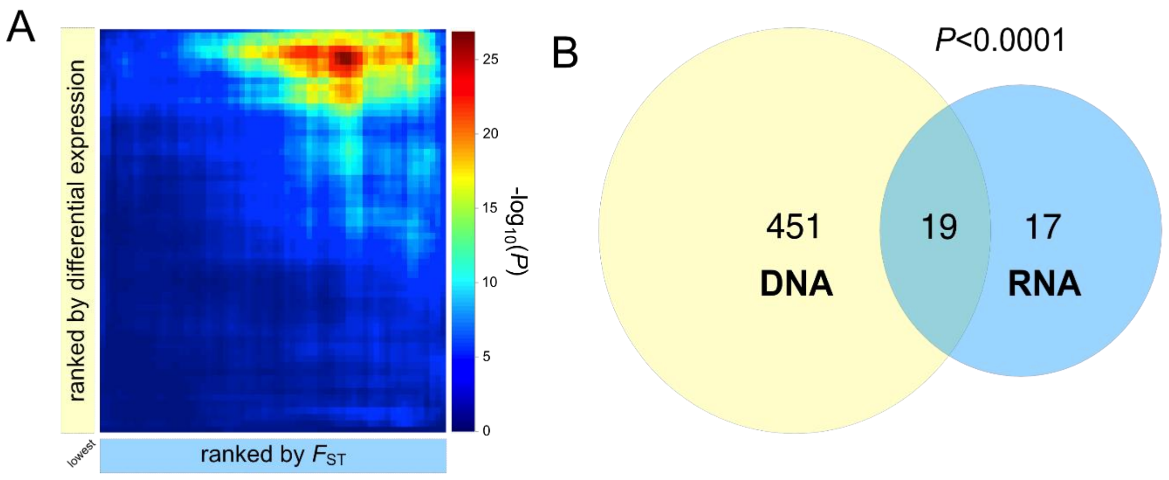


**Supplementary figure S9.** Overlaps between genomic and transcriptomic candidate genes (FIFS = Florida inverted vs. Florida standard) associated with *In(3R)Payne* independent of developmental temperature (similar to main text fig. 9 but with FIFS rather than FIFS_18 candidates)*.* (A) Summary of results of rank-rank hypergeometric overlaps. The dark red area indicates highly significant overlaps between genomic and transcriptomic candidates. (B) Venn diagram showing a highly significant overlap between genomic candidate genes (based on candidate SNPs with $F_{ST} \geq 0.9$ between karyotypes in Florida; in light yellow) and transcriptomic candidate genes (based on significant differential expression between karyotypes at 18°C; light blue); the *p*-value was estimated using the *SuperExactTest* package in *R*.

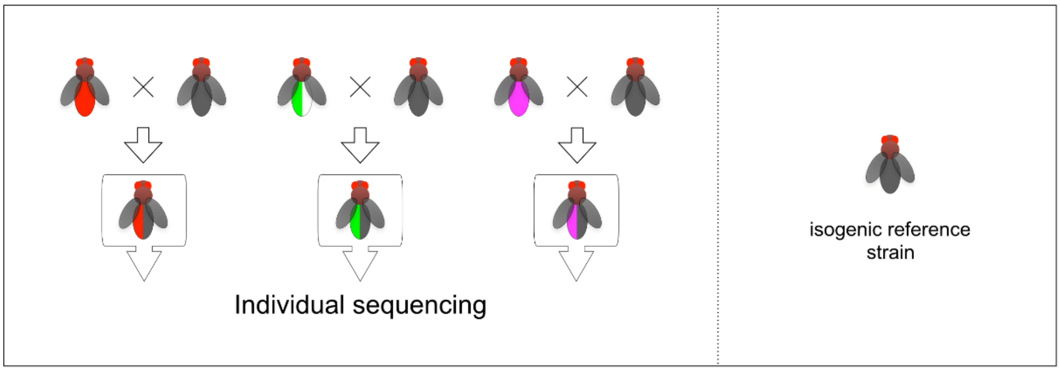


**Supplementary figure S10.** Crossing scheme depicting the generation of F1 hemiclones using a highly inbred reference strain, allowing to obtain high-confidence haploid genomic sequencing data through bioinformatic phasing. See Materials and Methods for details.
